# First-male sperm precedence in polyandrous *Spodoptera frugiperda* allows sterile males induce population suppression

**DOI:** 10.1101/2024.07.10.602994

**Authors:** Hao Sun, Ling-Ao Bu, Xin-Yue Zhang, Zhi-Ruo Zhang, Ling-Yi Zhu, Shao-Cong Su, Di Guo, Gao Hu, Cong-Fen Gao, Subba Reddy Palli, Jackson Champer, Shun-Fan Wu

## Abstract

Males respond to intense sperm competition by adapting reproductive strategies to promote fertilization success, which is critical for population reproduction. Thus, investigating the patterns and mechanisms of sperm competition is crucial for the development and application of pest population management techniques. In this study, we analyzed the sperm precedence pattern of a major pest, the fall armyworm, and used this pattern to manage the pest population. First, we found that females had a post-mating response and did not gain direct benefit through multiple mating. Next, in a double mating experiment, we used a molecular marker created by CRISPR/Cas9 to determine that most females use only the sperm of the first male to produce offspring. To further explore the role of fertilizing sperm in sperm competition, we constructed a sterile male line with eupyrene sperm defect by knocking out the *B2t* gene. Interestingly, two round mating assays showed that first mating with *B2t*-null males inhibited sperm fertilization from a second wild-type male. In other words, prior mating with *B2t*-null males significantly reduced the fertility and fecundity of females. Based on this finding, we continued to explore whether sperm-deficient sterile males could be used in the management of FAW populations. Cage experiments and mathematical modeling analyses showed that the release of excess *B2t*-null males induced population suppression. Our study expands our knowledge of sperm competition patterns in lepidopteran. In addition, our study provides a paradigm to develop and apply genetic control methods based on sperm competition outcome in polyandrous pests.

**Significance:** Sperm competition is essential for maintaining population reproduction. Understanding patterns and mechanisms of sperm competition facilitates the development of appropriate pest genetic control methods. Here, we describe that a globally major pest, the fall armyworm displays the first-male sperm precedence pattern. Interestingly, first mating with *B2t*-null males, which produces non-functional eupyrene sperm, significantly reduces the fertility and fecundity of females. That means that the ejaculate of the first male, even if its eupyrene sperm are defective, can inhibit sperm fertilization from a second wild-type male. Based on this, the release of excess *B2t*-null males significantly suppresses FAW populations. These results suggest that future development of genetic control techniques based on targeting nucleated sperm can effectively control FAW populations.

## Introduction

When we think of competition in the biological evolution of organisms, we often think of all kinds of fierce fights and competitions. However, there is one importance type of competition that is hidden in the microscopic world that plays a crucial role in the reproduction and evolution of species - sperm competition (1). Females can mate with more than one potential male in a reproductive cycle, which causes sperm from multiple males to compete for fertilization in the female reproductive tract (1). In internal fertilization, females may bias fertilization to produce offspring using sperm from specific males, which is often called female cryptic choice (2, 3). The reproductive fluids of females play crucial roles in regulating the outcome of sperm competition to suit their reproductive interests (4). Thus, paternity success outcome during postcopulatory sexual selection arises from a complex interaction between female-initiated cryptic selection and males engaging in sperm competition (4, 5).

Since Geoffrey Parker defined the concept of sperm competition (1), sperm competition as a critical mechanism of post-mating sexual selection has been rapidly studied, particularly with respect to its genetic basis (6, 7). *Drosophila melanogaster* was used as a powerful genetic tool to deeply resolve the mechanism of sperm competition (6, 7). In *D. melanogaster*, the second-male sperm preference pattern is attributed to sperm displacement and a seminal-fluid effect that causes the incapacitation of stored first male sperm (8, 9). Substantial amounts of seminal proteins are transferred with the sperm and may alter female physiological behaviors in independent or collaborative ways, such as post-mating responses, thereby increasing reproductive success (10). Currently, the importance of seminal proteins for paternity outcomes has been recognized in a wide range of species (11, 12). The effect of some seminal fluid protein genes on paternity share was learned using *Drosophila* as a model through population genetic association analysis and functional gene studies (6, 7). For example, the accessory gland-derived *Acp36DE* gene is required for sperm storage and mating plug formation, and deletion of this gene results in a reduced paternity share for males (13). Though the phenomenon of sperm competition has been extensively described (14, 15), we still lack an understanding of the link between sperm preference patterns (or paternity outcomes) and managing the reproduction in animal populations.

The fall armyworm (FAW), *Spodoptera frugiperda* (Lepidoptera, Noctuidae) originated in Central and South America and has invaded Africa, Asia, and Australia, becoming a major invasive pest around the globe (16, 17). It has a highly polyphagous caterpillar that can feed on more than 350 plant species, with corn as a dominant host (18). Adults exhibit high fecundity coupled with strong migratory ability. These biological traits give FAW robust environmental adaptability, which has led to its widespread distribution, causing significant economic losses worldwide (19). In the field, the primary means of controlling FAW are chemical insecticides and planting of corn engineered to produce *Bacillus thuringiensis* (Bt) proteins (16, 17). However, existing resistance that leads to decreasing efficacy of many chemical insecticide families and transgenic Bt corn against FAW has been documented in 250 cases (http://www. pesticideresistance.org). Recently developed genetic control methods such as CRISPR/Cas9-based precision guided SIT (pgSIT) and gene drive could be used as alternative strategies for control of this pest in the future (20, 21). Therefore, investigating the pattern and mechanism of sperm competition in FAW is crucial for the practical application of genetic control techniques.

In this study, we first investigated mating and post-mating behavior of FAW. Next, we explored whether multiple mating provides direct benefits to female adults. In a double-mating experimental design, we used molecular markers created by CRISPR/Cas9 technology to determine paternity share. To initially explore the mechanism behind sperm competition, we constructed sterile males with eupyrene sperm defect by knocking out the *B2t* gene. Combined with double mating assays, we explored the effects of eupyrene sperm on the paternity outcome. Intriguingly, first mating with *B2t*-null males inhibited sperm fertilization from second WT males. Based on the natural reproductive strategy of FAW, we conducted cage experiments and mathematical modeling to demonstrate that releasing excess *B2t*-null males can significantly suppress FAW populations.

## Results

### FAW displays post-mating response and multiple mating behavior

We first studied the characteristics of post-mating response and multiple mating behavior in females. Within 48 hours after the first mating, the remating rate of gravid females decreased significantly (**Figure 1A**). The mated females also gained a dramatic reproductive advantage. The mated females laid significantly higher numbers of eggs than the virgin females (**Figure 1B**). We investigated the distribution of the number of mating times of females in the laboratory. The study showed that about 73% of the adult females were receptive to multiple mating (**Figure 1C**). A few females (<10%) mated up to 5 or 6 times (**Figure 1C**).

**Figure 1.**
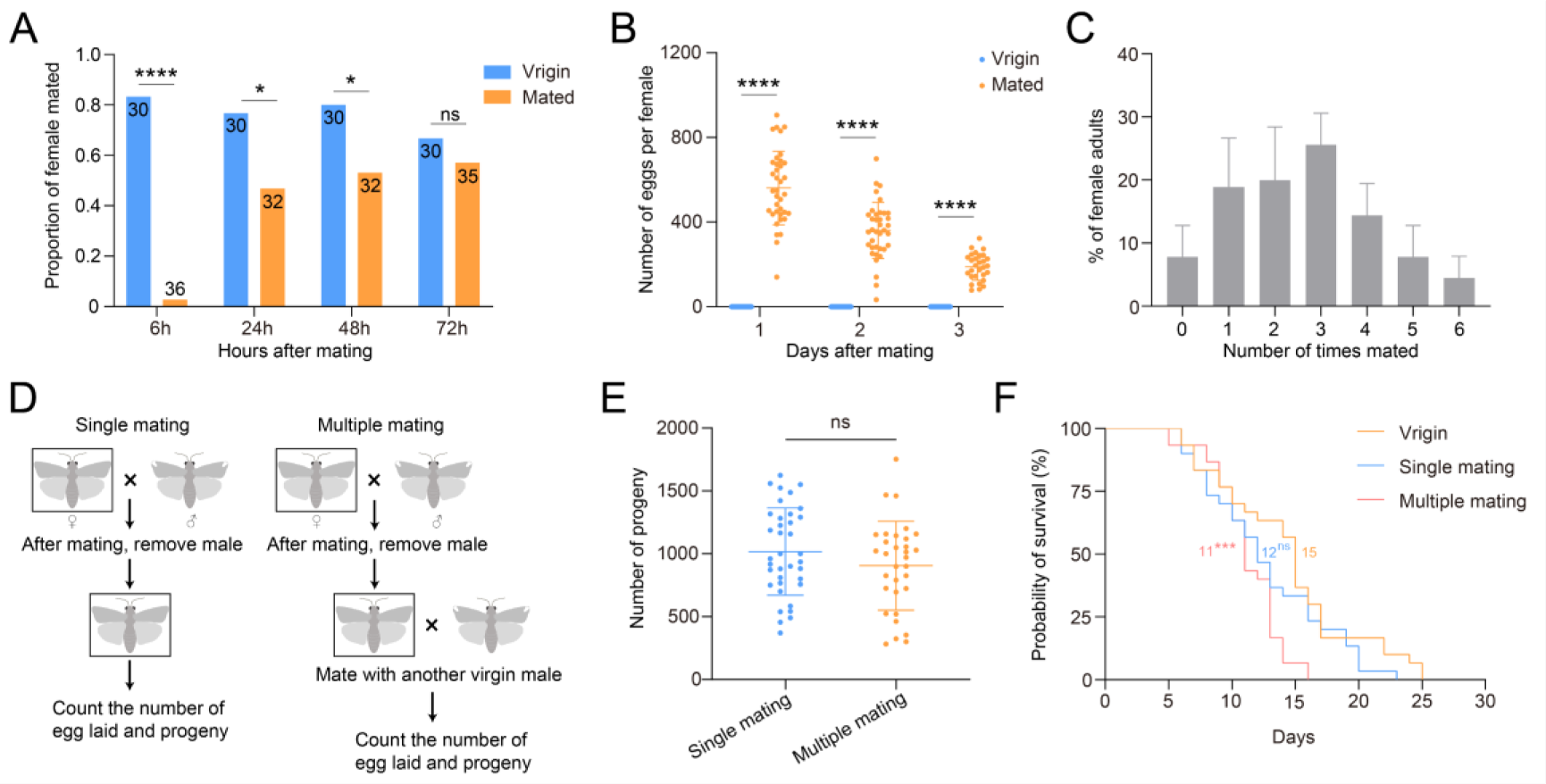
FAW had a post-mating behavior and did not receive direct benefits from the multiple mating. (**A**) Re-mating rate of females at 6, 24, 48 and 72h after first mating. Fisher’s exact test; ns, not significant, \**P* < 0.05, \*\*\*\**P* < 0.0001. (**B**) Quantification of eggs laid by females on 1-, 2-, and 3-days post-mating. n = 22 - 37. One-sample Student’s t test versus zero; \*\*\*\**P* < 0.0001. (**C**) Proportion of estimated mating times per adult female. (**D**) Schematic of the single and multiple mating of females. (**E**) The number of progeny produced by each female after single and multiple mating. n = 32 - 37. Student’s t test; ns, not significant. (**F**) Comparison of adult female longevity in virgins, single and multiple mating states. n = 30. Statistics were performed using the Log-rank (Mantel-Cox) test; ns, not significant, \*\*\**P* < 0.001. The median survival number is displayed on the picture.

We then investigated the potential benefits of multiple mating for females. Schematic diagrams of single and multiple mating experimental designs for females are shown in **Figure 1D**. There were no significant differences in egg production and egg hatchability between females mated multiple (2.7 times on average) or single time (**Figure 1E, Figure 1-figure supplement 1A-B**). Females produced the highest number of eggs on the first day after mating, with a significant decrease on the second and third days (**Figure 1-figure supplement 1C**). In contrast, the longevity of females that mated multiple times was significantly lower than that of once-mated and virgin females. The longevity of polyandrous females was significantly lower than that of once-mated and virgin females (**Figure 1F**). These results suggested that females did not gain direct benefits from multiple mating.

Meanwhile, we explored the effects of second mating on fertility, mating behavior, and spermatophore formation in males through two rounds of single-pair mating trials (**Figure 1-figure supplement 2A**). Within 6 hr after the first mating, the remating ability of mated males was significantly reduced (Virgin, 87.14%; 6 hr after first mating, 4.92%). In contrast, at 24 h after mating, the remating rate of mated males did not differ from the first mating rate (Virgin, 87.14%; 24h after first mating, 84.00%; *P* = 0.7914) (**Figure 1-figure supplement 2B**). In addition, multiple mating had no significant effect on male mating duration and fertility (**Figure 1-figure supplement 2C-E**). A gradual decrease in the daily egg production of females was recorded over a three-day post-mating period (**Figure 1-figure supplement 2F**). Immediately after mating, spermatophore was isolated from the copulatory bursa of females. No difference in the frontal cross-sectional area of the spermatophore formed was detected after the first and second mating (**Figure 1-figure supplement 2I**). Interestingly, the color of the spermatophore was different in the two cases (**Figure 1-figure supplement 2G-H**). Whether this difference in coloration is related to the fact that males allocate seminal fluid with different functions depending on the number of copulations or position in the mating order deserves further study.

**Figure 1-figure supplement 1.**
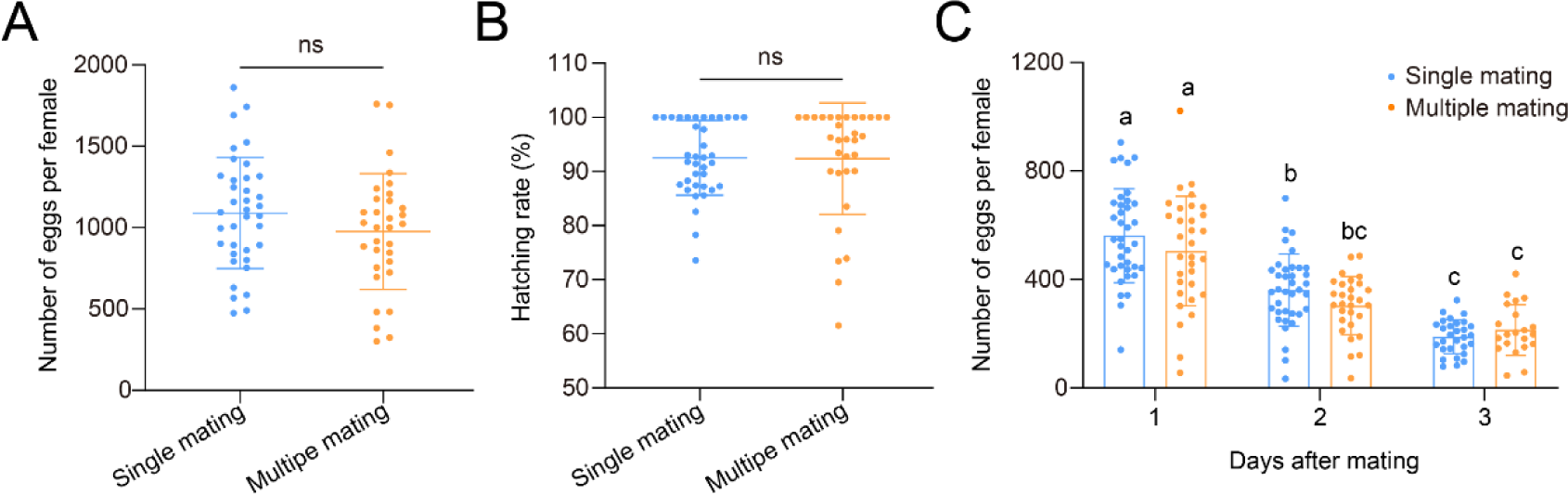
The effects of multiple mating on female egg-laying behavior. (**A**) The number of eggs laid by each female after single and multiple mating. n = 32 - 37. Student’s t test; ns, not significant. (**B**) The hatch rate (%, the number of larvae hatched divided by the total number of eggs laid) of eggs laid by females after single and multiple mating. n = 32 - 37. Mann–Whitney test; ns, not significant. (**C**) The number of eggs laid by females on three consecutive days after single or multiple mating. n = 21 - 37. One-way ANOVA followed by Tukey’s multiple comparisons test. The lowercase letters above each bar indicate significant differences (*P* < 0.05). All data are presented as means ± SD.

**Figure 1-figure supplement 2.**
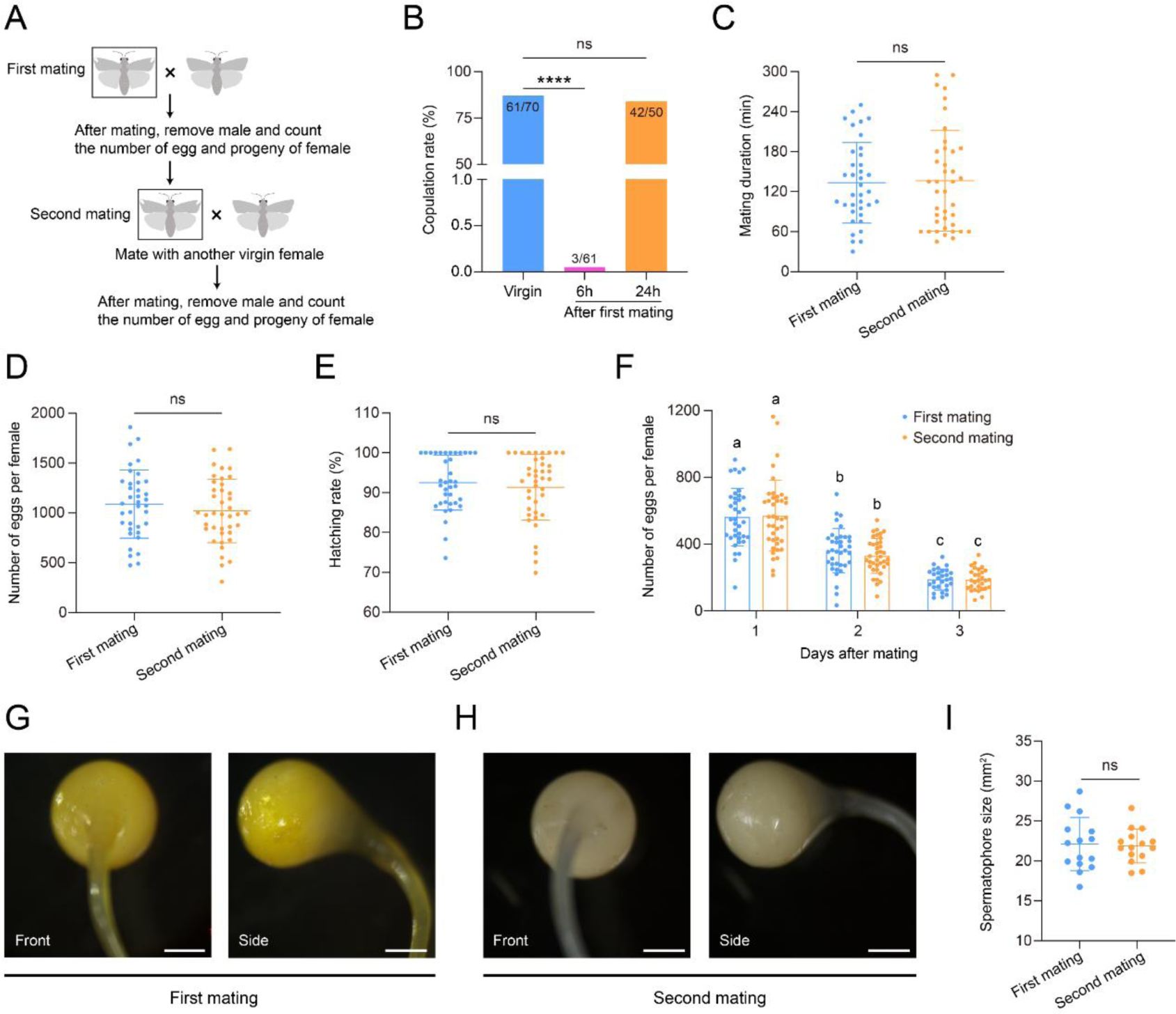
The effects of multiple mating on the reproductive behavior of males. (**A**) Schematic of the first and second mating of males. (**B**) After the first mating of males, the remating rate decreased significantly at 6 hr after mating, and the re-mating rate recovered to the level of the first mating rate at 24 hr after mating. Fisher’s exact test; ns, not significant, \*\*\*\**P* < 0.0001. (**C**) The mating duration of the first and second mating in males. n = 37 - 40. Student’s t test; ns, not significant. (**D**) The number of eggs per female after single or multiple mating. n = 37 - 40. Student’s t test; ns, not significant. (**E**) The hatching rate of eggs laid by females after single or multiple mating. The hatching rate (%) = the number of larvae hatched / the total number of eggs laid. n = 37 - 40. Mann–Whitney test; ns, not significant. (**F**) Eggs laid by females for three consecutive days after single or multiple mating. n = 29 - 40. Kruskal–Wallis test with Dunn’s multiple comparisons test. The lowercase letters indicate significant differences (*P* < 0.01). (**G and H**) Images of the front and side of the spermatophore after single (G) or multiple mating (H). Scare bar, 2 mm. (**I**) The cross-sectional area of the front of the spermatophore v. n = 15. Student’s t test; ns, not significant. All data are presented as means ± SD.

### First-male sperm precedence patterns present in FAW

Females mate with more than one male, which may result in the production of offspring of mixed paternity. Molecular markers such as microsatellite DNA are often used to determine the parental share of fertilized eggs (22). To study the sperm competition patterns of individual FAW, we designed a double mating experiment with different mating intervals (**Figure 2A**). We first used CRISPR/Cas9 gene editing to knock out the *NinaB* gene to create a molecular marker. A mixed system of Cas9 and gRNA was injected into fresh embryos, and after 3 days, 36.0% of larvae successfully hatched (**Table S2**). PCR identified 58.9% of the G0 adults as *NinaB* chimeric mutants (**Table S2**). After the genetic cross, we obtained *NinaB* homozygous mutants (*NinaB^-/-^*) with an overall deletion of 5 bp at the sgRNA site (**Figure 2-figure supplement 1**). Fecundity measurements indicated that *NinaB* mutation did not result in reduced fertility in males (**Figure 2-figure supplement 2**). After females second mated with *NinaB^-/-^* males, we identified the genotypes of the offspring produced after the second mating using F/R primers. Our study found that 79.2%, 81.8%, or 70.8% of females produced only WT male offspring at mating intervals of 1, 2, or, 3 days, respectively (**Figure 2B**). The results suggest that FAW individuals use a first-male sperm precedence pattern. In addition, the double mating test also demonstrated that females can gain genetic benefit through multiple mating, which favors genetic diversity in the offspring.

**Figure 2.**
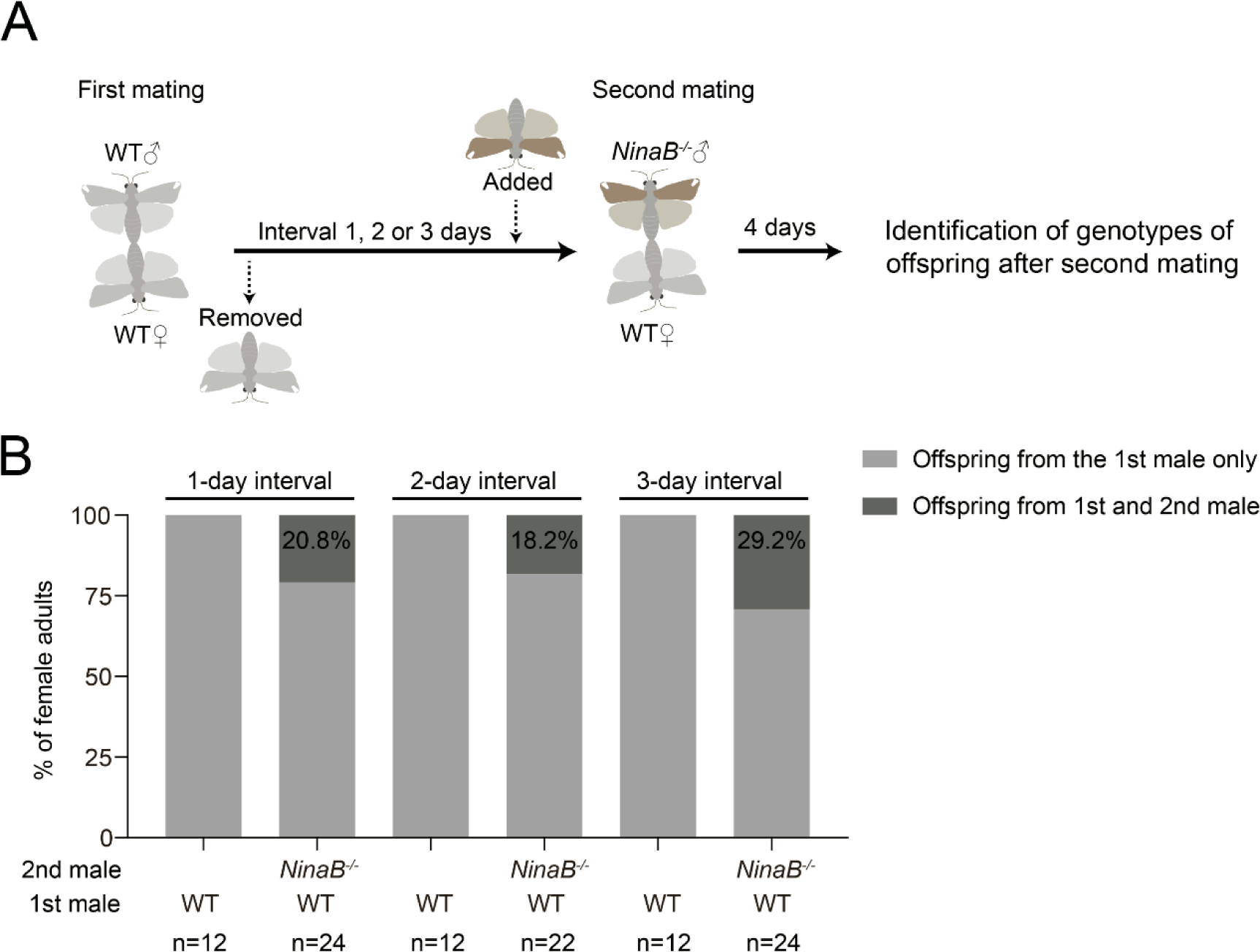
Most females produced offspring that were sired by the first male. (**A**) Schematic diagram of the double mating experiments. (**B**) FAW displayed first-male sperm precedence pattern in different mating intervals.

**Figure 2-figure supplement 1.**
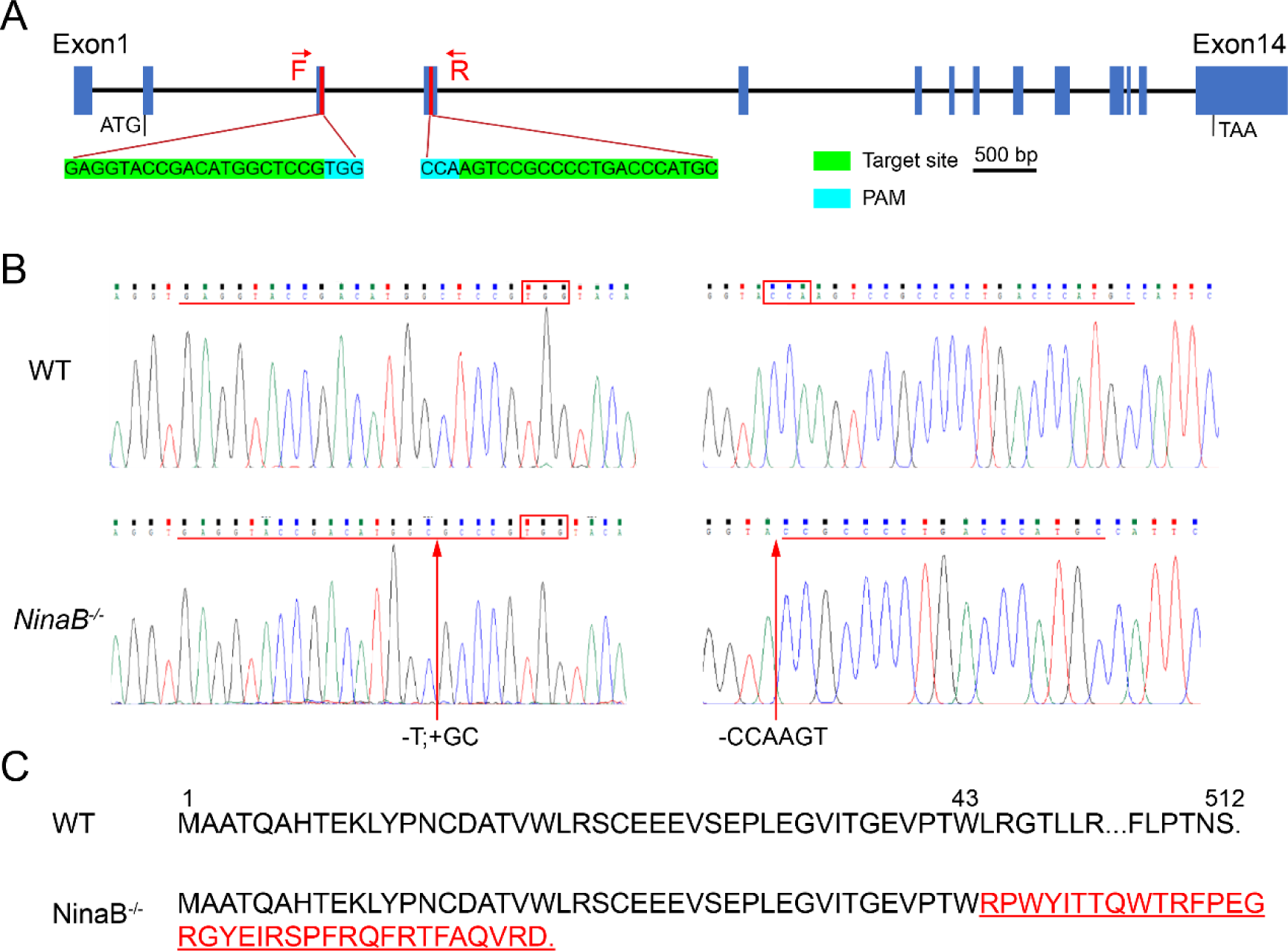
Disruption of *NinaB* gene using CRISPR/Cas9. (**A**) Design two gRNAs to target the *NinaB* gene. The *NinaB* gene includes 14 exons. Two gRNAs were designed to target exon 3 and exon 4, respectively. PAM sequences were highlighted in blue and target sequences were labeled in green. (**B**) The Sanger sequence chromatograms flanking the sgRNA target site for wild type (WT) and homozygous mutant (*NinaB^-/-^*). The sgRNA target sequence and PAM are underlined in red. (**C**) The predicted NinaB protein sequence of WT and *NinaB^-/-^*. A 7 bp deletion and 2 bp insertion of the mutant allele generated a truncated 78-residue protein.

**Figure 2-figure supplement 2.**
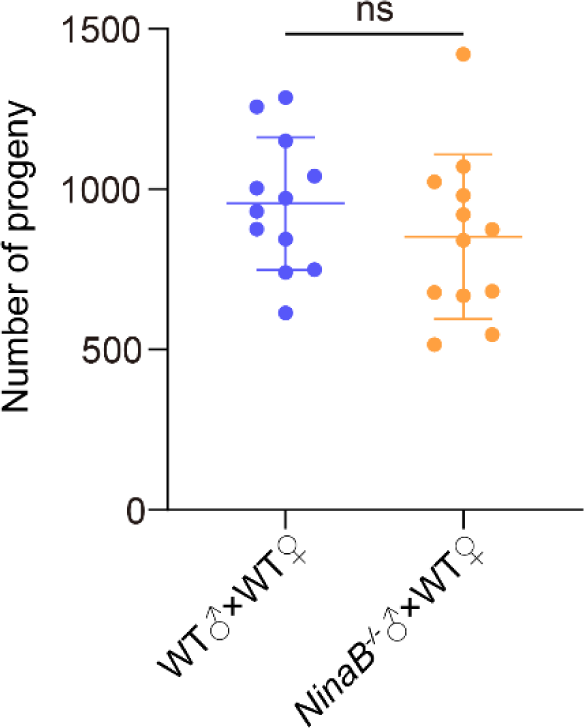
Determination of fertility in *NinaB^-/-^* males. Number of offspring resulting from crosses of WT males or *NinaB^-/-^*males with WT females. n = 12. Error bars are mean ± SD. Student’s t test; ns, not significant.

### *B2t* mutation causes male sterility

Previous studies showed that the testis-specific expression of the *B2t* gene is involved in spermatogenesis and is critical for male fertility in mosquito and *Drosophila* (23, 24). To obtain sperm-deficient sterile males, we employed CRISPR/Cas9 technology to knock out the *B2t* gene. The full-length *B2t* gene ORF was cloned from *S. frugiperda*, which consists of two exons and encodes 443 amino acids (**Figure 3A**). Amino acid sequence alignments showed that the *S. frugiperda* B2t protein is similar to its orthologs from other species (>95% identity) and also contains an EGEFXXX (X is an acidic residue) motif at the C-terminus, as proposed by Kawasaki (**Figure 3A-figure supplement 1A**). The phylogenetic analysis further revealed the *S. frugiperda* B2t clustered with the *B. mori* B2t, suggesting a possible conserved function in lepidopteran insects (**Figure 3A-figure supplement 1B**). RT-PCR and qRT-PCR were performed to examine the expression of *B2t* in adult tissues and throughout development, respectively. The results showed that the expression of *B2t* was limited to the male testis (**Figure 3A-figure supplement 2**). The ribonucleoprotein complex was injected into fresh eggs of *S. frugiperda*. Testing of progeny showed that 90.0% (36/40) of the tested G0 contained *B2t* locus mosaic mutations (**Table S2**). Subsequently, through multi-generational crosses, we successfully generated a homozygous mutant *B2t*^-/-^ with a 7-bp deletion (**Figure 3A-B, Figure 3A-figure supplement 3A-C**). The 7-bp deletion of *B2t* coded for a truncated 200-residue protein (**Figure 3A-figure supplement 3D**). qRT-PCR analysis showed that the expression of *B2t* is undetectable in *B2t*^-/-^ mutants (**Figure 3C**).

**Figure 3.**
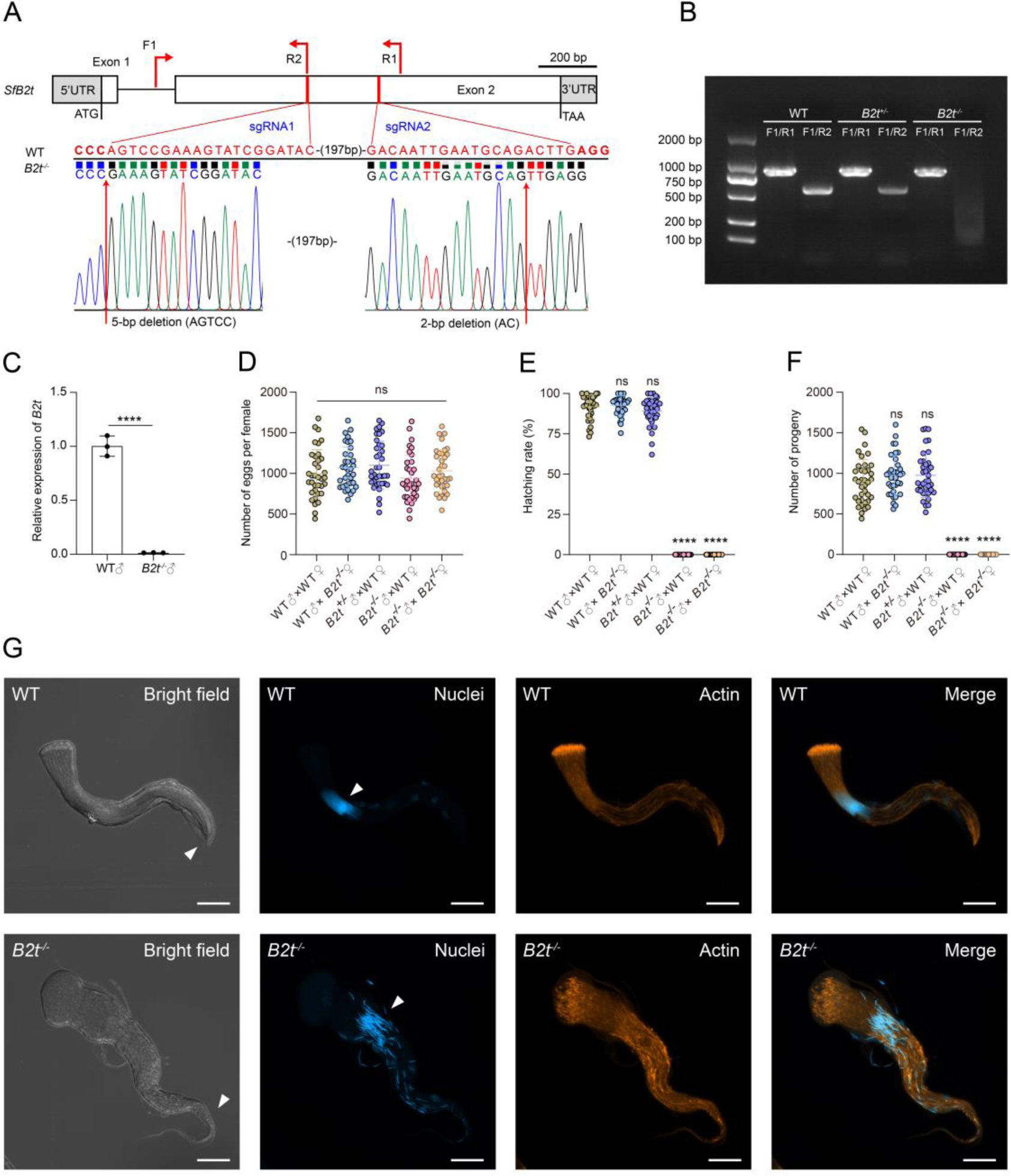
Disruption of *B2t* causes male sterility. (**A**) CRISPR/Cas9-based knockout of *B2t*. *B2t* consists of two exons. ATG, TAA, 5’ UTR and 3’ UTR are marked. Scale bar, 200 bp. Two single-guide RNAs (sgRNAs) were designed to target exon 2 of *B2t*. The sgRNA sequences are shown in red and the protospacer adjacent motif (PAM) sequences are shown in bold red. The positions of the primers (F1, R1, R2) used in (B) are indicated by red arrows. The *B2t* mutation with a 5-bp deletion at sgRNA1 (AGTCC) and a 2-bp deletion at sgRNA2 (AC) was confirmed by sequence chromatograms. (**B**) Genotyping of *B2t* mutation. The primer pair F1/R1 was used to check if *B2t* was mutated. The homozygous *B2t* mutants (*B2t^-/-^*) were identified by the primer pair F1/R2. (**C**) The mRNA expression level of *B2t* in the testis of wildtype (WT) and *B2t^-/-^* adults. n = 3. Student’s t test; \*\*\*\**P* < 0.0001. (**D**) The number of eggs per female. n = 34 - 37 for each treatment. One-way ANOVA followed by Tukey’s multiple comparisons test; ns, not significant. (**E**) The hatch rate [(%) = the number of hatched larvae/the number of eggs laid] by WT or mutant females crossed with WT or mutant males. n = 34 - 37 for each treatment. Kruskal–Wallis test with Dunn’s multiple comparisons test; ns, not significant, \*\*\*\**P* < 0.0001. (**F**) The number of progeny produced per mating combination. n = 34 - 37 for each treatment. Kruskal–Wallis test with Dunn’s multiple comparisons test; ns, not significant, \*\*\*\**P* < 0.0001. (**G**) *B2t* mutation interferes with the development of eupyrene sperm bundles. Representative immunofluorescence images of eupyrene sperm bundles from WT males and *B2t^-/-^* males, showing abnormal tail morphology of sperm bundles and location and spacing of sperm nuclei caused by *B2t* knockout. White triangular symbols show the differences. The nuclei (Blue) were stained with DAPI. The filamentous actin proteins (Red) were stained with TRITC Phalloidin. Scale bar, 20 μm. All data are presented as means ± SD.

To test whether disruption of *B2t* impairs male fertility, we conducted single-pair mating assays in 475 mL plastic cups. Five treatments were used in this assay: WT males + WT females, WT males + *B2t*^-/-^ females, *B2t*^+/-^ males+ WT females, *B2t*^-/-^ males+ WT females, and *B2t*^-/-^ males + *B2t*^-/-^ females. The fecundity measurement test showed that disruption of *B2t* leads to male sterility (**Figure 3D-E**). We then explored the mechanism behind male sterility caused by *B2t* mutation. Imaging of eupyrene sperm bundles showed that the sperm bundles tail of *B2t^-/-^* males is abnormal (**Figure 3G**). The sperm nuclei of *B2t^-/-^* males were scattered and disordered compared to the neat and tightly arranged sperm nuclei of WT males (**Figure 3G**). However, deletion of *B2t* did not affect the development of apyrene sperm (**Figure 3A-figure supplement 4**). We recorded the development of eggs laid by WT females mated with *B2t^-/-^* males. Eggs from cross of WT female with *B2t^-/-^* males showed no obvious signs of embryonic development compared to WT eggs (**Figure 3A-figure supplement 5A**). Genotyping of unhatched eggs showed that these eggs did not inherit the *B2t* mutant allele, which could mean that *B2t* deficient sperm could not fertilize eggs. (**Figure 3A-figure supplement 5B**).

**Figure 3-supplement 1.**
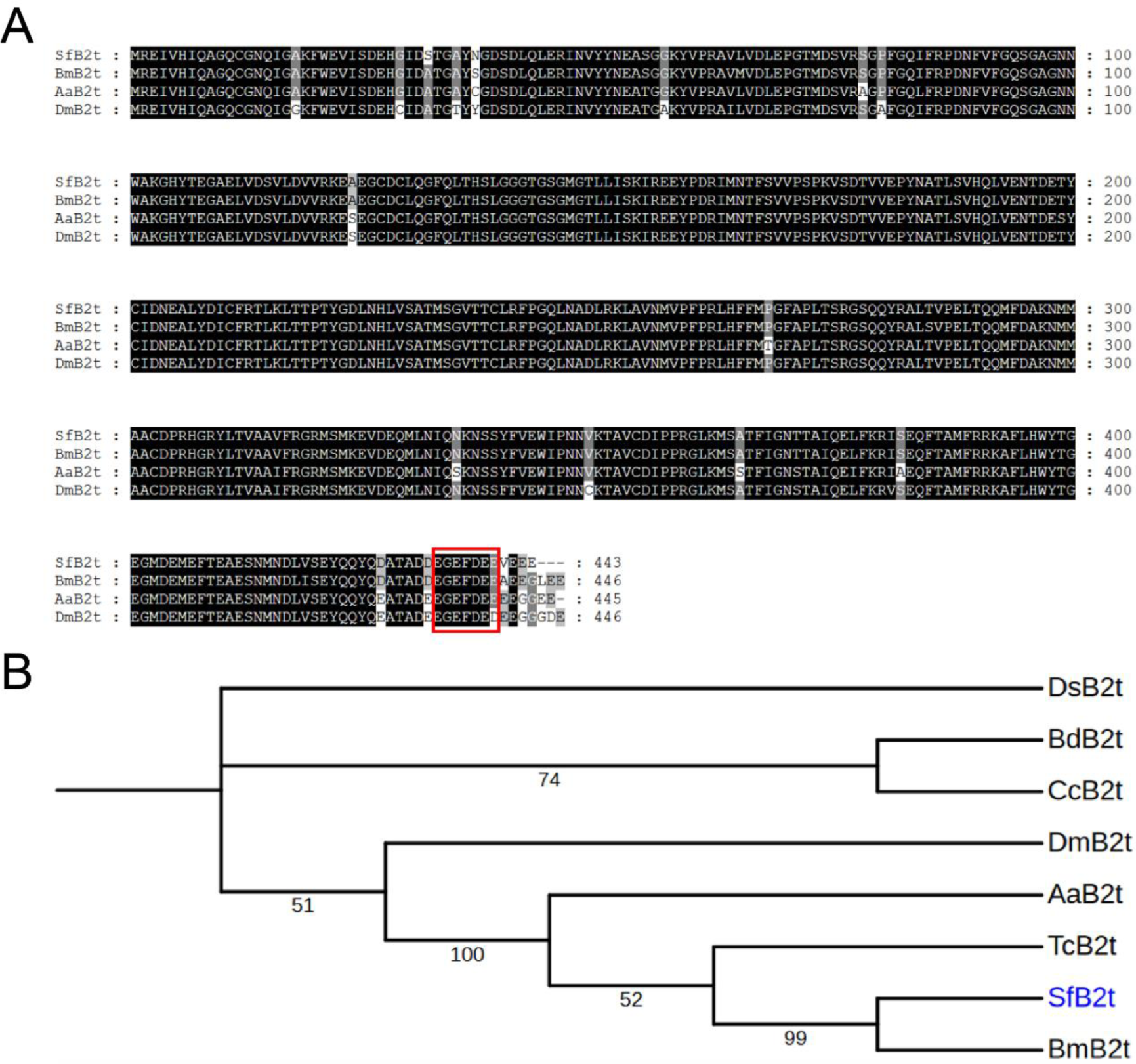
Molecular characterization of B2t. (**A**) Amino acid sequence alignment of B2t protein. The C-terminal axoneme motif EGEFXXX (X is an acidic residue) is indicated in the red box. Protein sequence of *Bombyx mori* B2t (BmB2t, NP_001037060.1), *Aedes aegypti* B2t (AaB2t, XP_030036707.1), and *Drosophila melanogaster* B2t (DmB2t, AAT71306.1) were obtained from the GenBank database. (**B**) Phylogenetic analysis of *S. frugiperda* B2t (SfB2t) and its orthologs from other insects. The phylogenetic tree was constructed by MEGA-X using a neighbor-joining method. Bootstrap values were calculated after 1000 replicates and displayed on each node. Abbreviations and GenBank accession numbers as follow: DsB2t: *Drosophila suzukii* B2t (XP_016937909.1), BdB2t: *Bactrocera dorsalis* B2t (JAC41313), CcB2t: *Ceratitis capitata* B2t (EU386342), DmB2t: *Drosophila melanogaster* B2t (NP_524290), AaB2t: *Aedes aegypti* B2t (DQ833526), TcB2t: *Tribolium castaneum* B2t (ACB45871.1), and BmB2t: *Bombyx mori* B2t (CAA52906).

**Figure 3-supplement 2.**
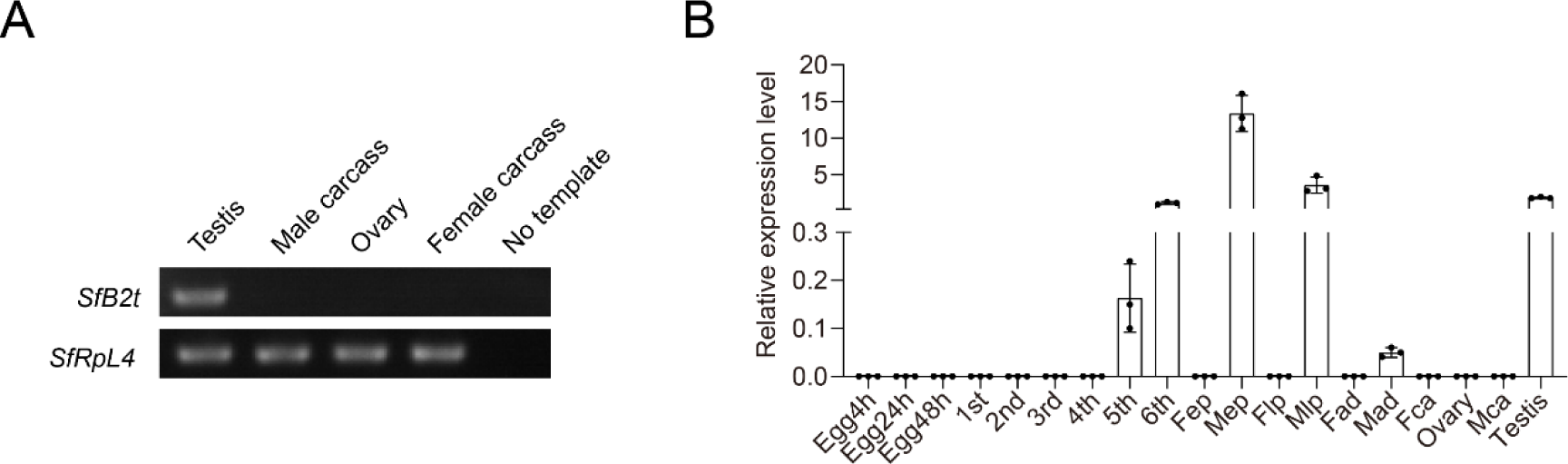
*B2t* is specifically expressed in the testis. (**A**) RT-PCR analysis of the expression of *B2t* mRNA in adult tissues. The housekeeping gene *RpL4* was used as an internal control. (**B**) The spatiotemporal expression profile of *B2t* quantified by qRT-PCR. The mean ± SD from three biological replicates. Abbreviations for samples: 1st: First instar larva; 2nd: Second instar larva; 3rd: Third instar larva; 4th: Fourth instar larva; 5th: Fifth instar larva; 6th: Sixth instar larva; Fep: Female early pupa; Mep: Male early pupa; Flp: Female late pupa; Mlp: Male late pupa; Fad: Female adults; Mad: Male adults; Fca: Female carcass (Female adult without ovary); Mca: Male carcass (Male adult without testis).

**Figure 3-supplement 3.**
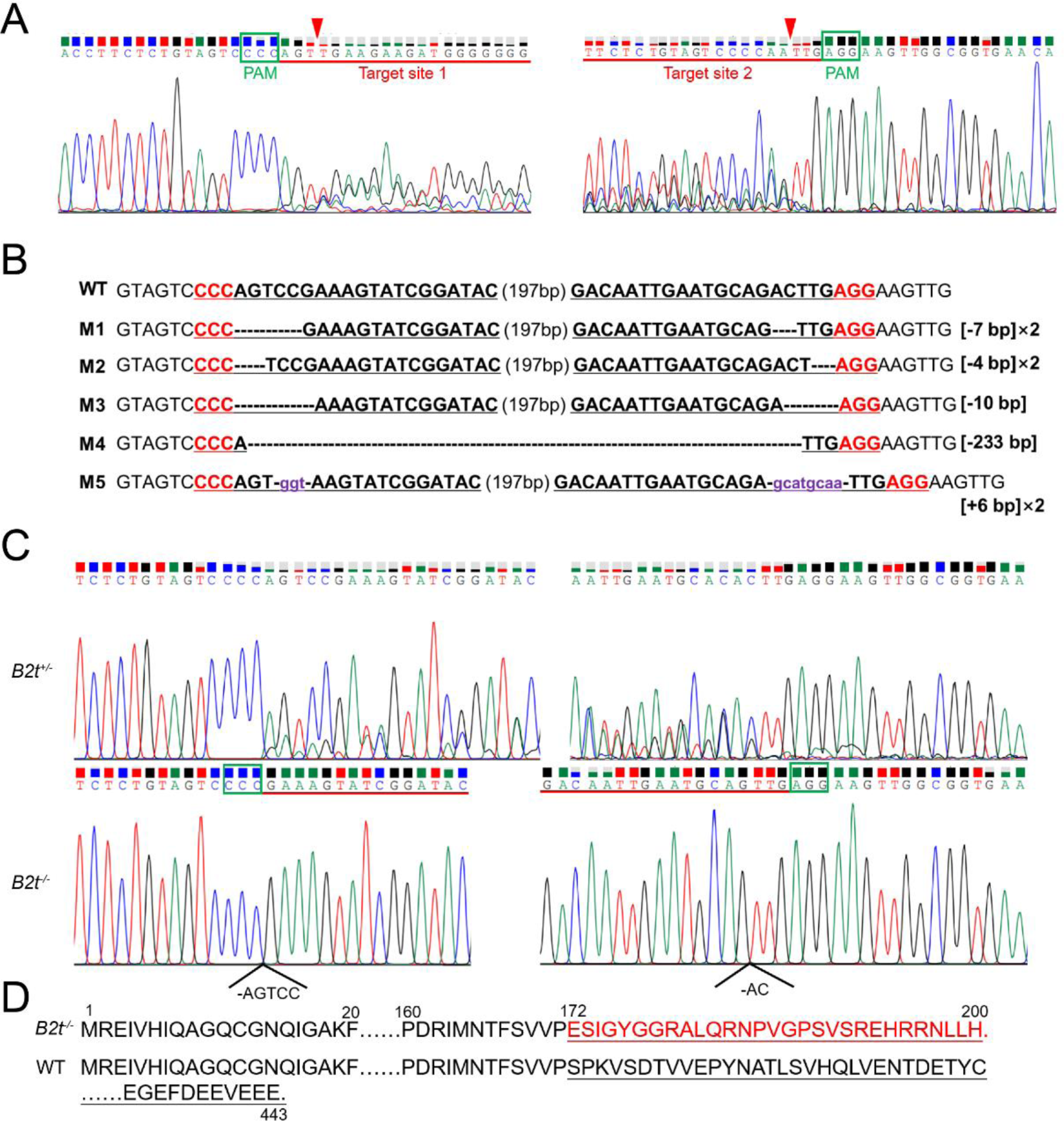
*B2t* mutation analysis. (**A**) Representative chromatogram of direct sequencing of G0 individuals PCR products indicating two sgRNA sites were successfully cleaved. The two sgRNA sequences are underlined in red, and a green box indicates the PAM sequence. The position of the red triangle indicates the Cas9 protein cleavage site. (**B**) TA cloning and sequencing showed the insertion/deletion (indel) in the target site. Two sgRNAs (bold black) and PAM sequence (bold red) are underlined. In the mutated sequence (M), the dashed lines indicate the deletion, and the blue lowercase letters are the inserted sequences. The change in the sequence length [+, insertion; -, deletion]. ×n indicates that the mutant was detected in n sequenced clones. (**C**) The sequence chromatograms flanking the target site for heterozygous mutant (*B2t^+/-^*), and homozygous mutant (*B2t^-/-^*) with a 7 bp deletion. Red lines indicate the sgRNA and green boxes indicate PAM sites. (**D**) Comparison of protein sequences of WT and *B2t^- /-^* mutant. Numbers denote the length of the protein sequences. Miscoded protein sequences are indicated by red letters in *B2t^-/-^* mutant.

**Figure 3-supplement 4.**
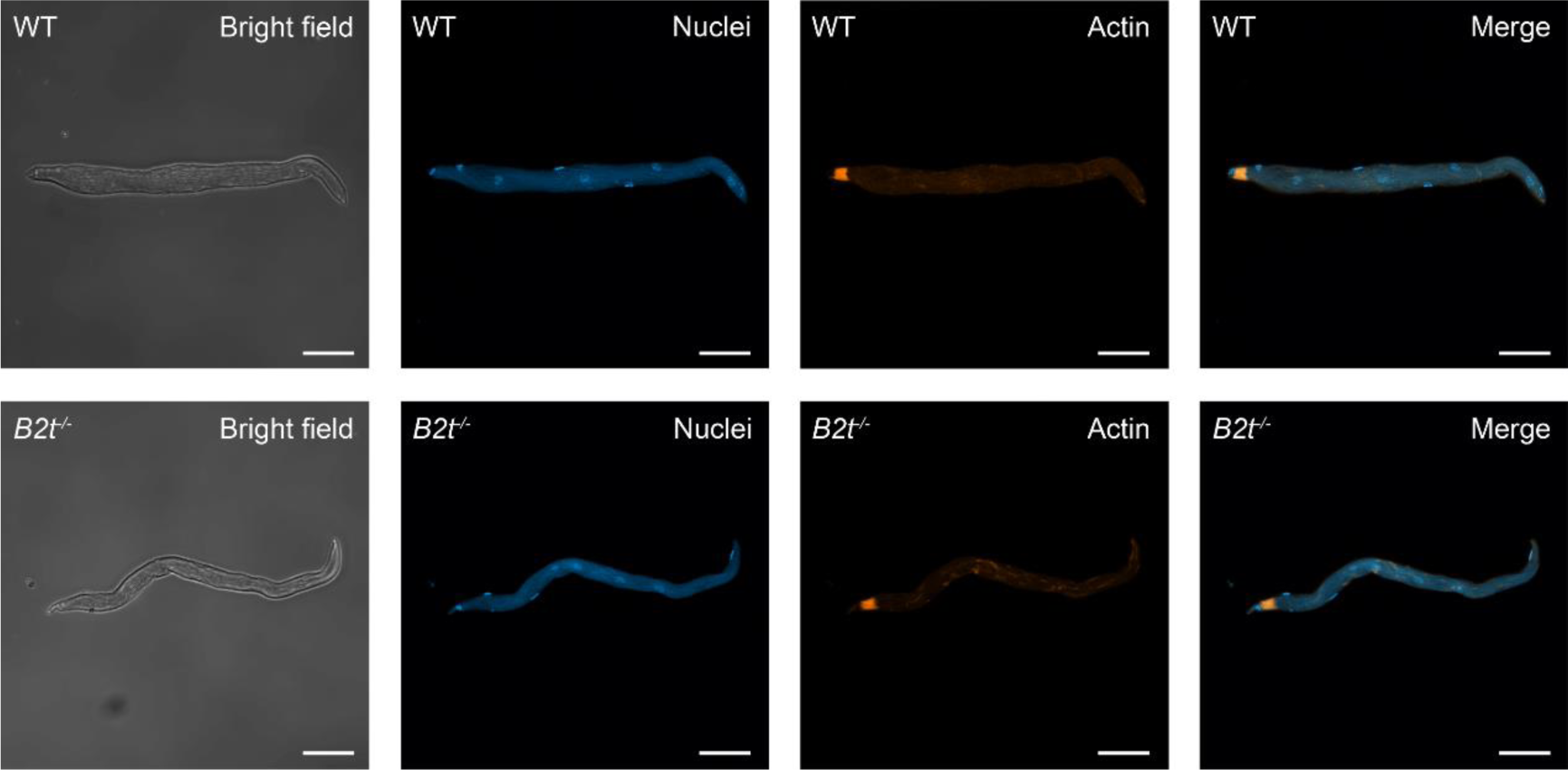
Disruption of *B2t* does not affect the development and morphology of apyrene sperm bundles. The nuclei (Blue) were stained with DAPI. The filamentous actin proteins (Red) were stained with TRITC Phalloidin. Scale bar, 20 μm.

**Figure 3-supplement 5.**
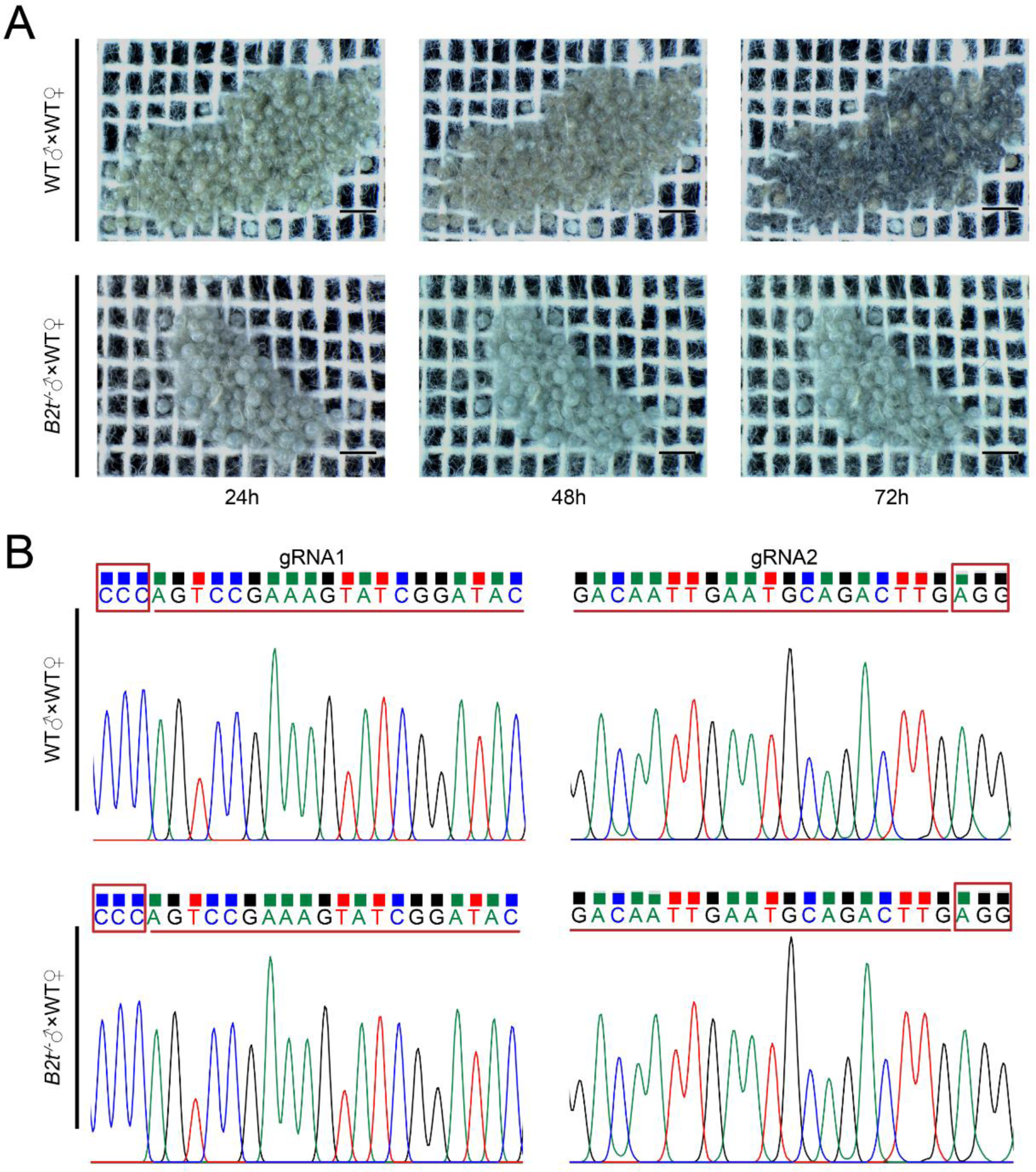
*B2t*-deficient eupyrene sperm could not fertilize eggs. (**A**) Photographing the developmental state of eggs at three time points (24h, 48h and 72h after spawning). Eggs were laid by WT males + WT females and *B2t*^-/-^ males+ WT females. Scale bar, 1000 μm. (**B**) Direct sequencing chromatogram of PCR products used to determine the genotype of eggs in (A). The gRNA sequence is underlined in red, and the PAM sequence is in the red box.

### Prior mating with *B2t*^-/-^ males inhibited sperm fertilization from a second WT male

The results of competitive fertilization showed that most of the offspring produced after the second mating were from the first male only. To reveal the mechanism of first-male sperm precedence, we performed two rounds of mating experiments with different mating intervals. Females were mated for the first time with *B2t*^-/-^ males that produced non-functional eupyrene sperm, and after an interval of 1, 2, or 3 days, they were mated again with WT males (**Figure 4A**). Within 24 h after mating with *B2t*^-/-^ males, females had a postcopulatory response, with a significant decrease in the rate of mating again with WT males (**Figure 4B**). Interestingly, we found that prior mating with *B2t*^-/-^ males could inhibit fertilization of sperm from WT second males at different mating intervals (**Figure 4C-E**). Overall results showed that after both rounds of mating, 75% of females would still not produce viable offspring (**Figure 4F**). Although 25% females were fertile, their fecundity was significantly suppressed (**Figure 4G**). Population genetic analysis studies suggested that apyrene sperm in Lepidoptera do not directly affect the outcome of sperm competition (25). Combined with our studies, we conclude that seminal fluid proteins likely play a key role in the first-male sperm precedence. We hypothesize that seminal fluid proteins may block the sperm of the second male from entering the spermatheca and binding to oocytes in an independent or synergistic manner.

**Figure 4.**
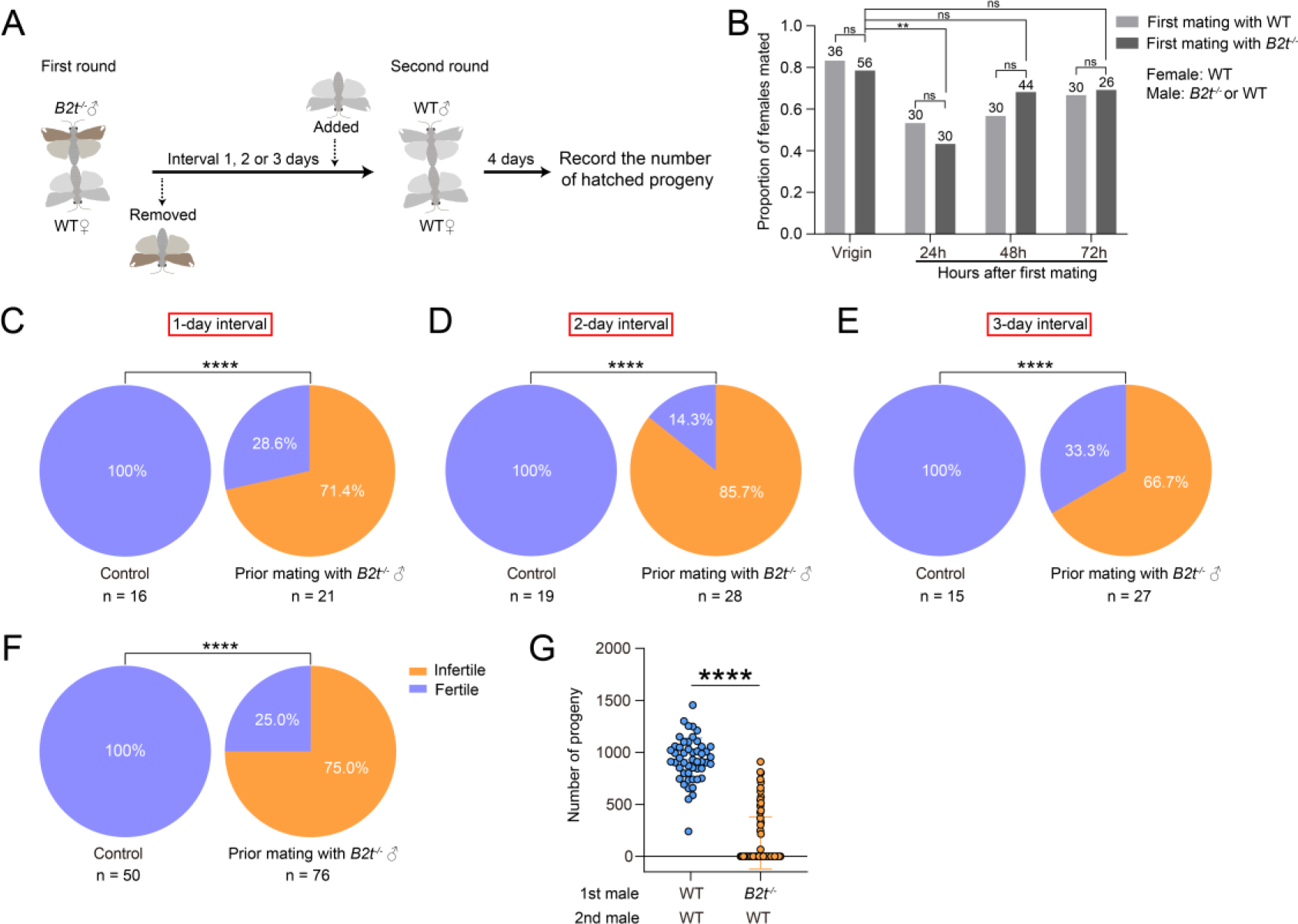
The effect of the *B2t* mutation on paternity. (**A**) Schematic diagram of a double mating test with different mating intervals. (**B**) Proportion of WT females re-mating with virgin WT males after first mating with WT or *B2t^-/-^* males. Fisher’s exact test; ns, not significant; \*\**P* < 0.01. Female: WT; Male: *B2t^-/-^* or WT. (**C**) Percentage of fertile females in double mating trials at 1-day intervals. Fisher’s exact test; \*\*\*\**P* < 0.0001. (**D**) Percentage of fertile females in double mating trials at 2-day intervals. Fisher’s exact test; \*\*\*\**P* < 0.0001. (**E**) Percentage of fertile females in double mating trials at 3-day intervals. Fisher’s exact test; \*\*\*\**P* < 0.0001. (**F**) Percentage of fertile females in double mating trials with different days of interval. Fisher’s exact test; \*\*\*\**P* < 0.0001. (**G**) Prior mating with *B2t^-/-^* males inhibit female fertility. The number of progeny from the double mating assay. n = 50 - 76. Error bars are mean ± SD. Student’s t test; \*\*\*\**P* < 0.0001.

In addition, we asked whether a second mating with *B2t*^-/-^ males would reduce the number offspring produced by females when the first male was WT. The experimental procedure was the same as described above, but the mating order of the males was reversed (**Figure 4-figure supplement 4A**). The results showed that the fertility and fecundity of females first mated with WT males would not be reduced at any of our mating intervals. (**Figure 4-figure supplement 4B-C**). Compared to the results of first round mating with *B2t*^-/-^ males, we found that *B2t*-null males were disadvantaged in some aspects of sperm competition.

**Figure 4-supplement.**
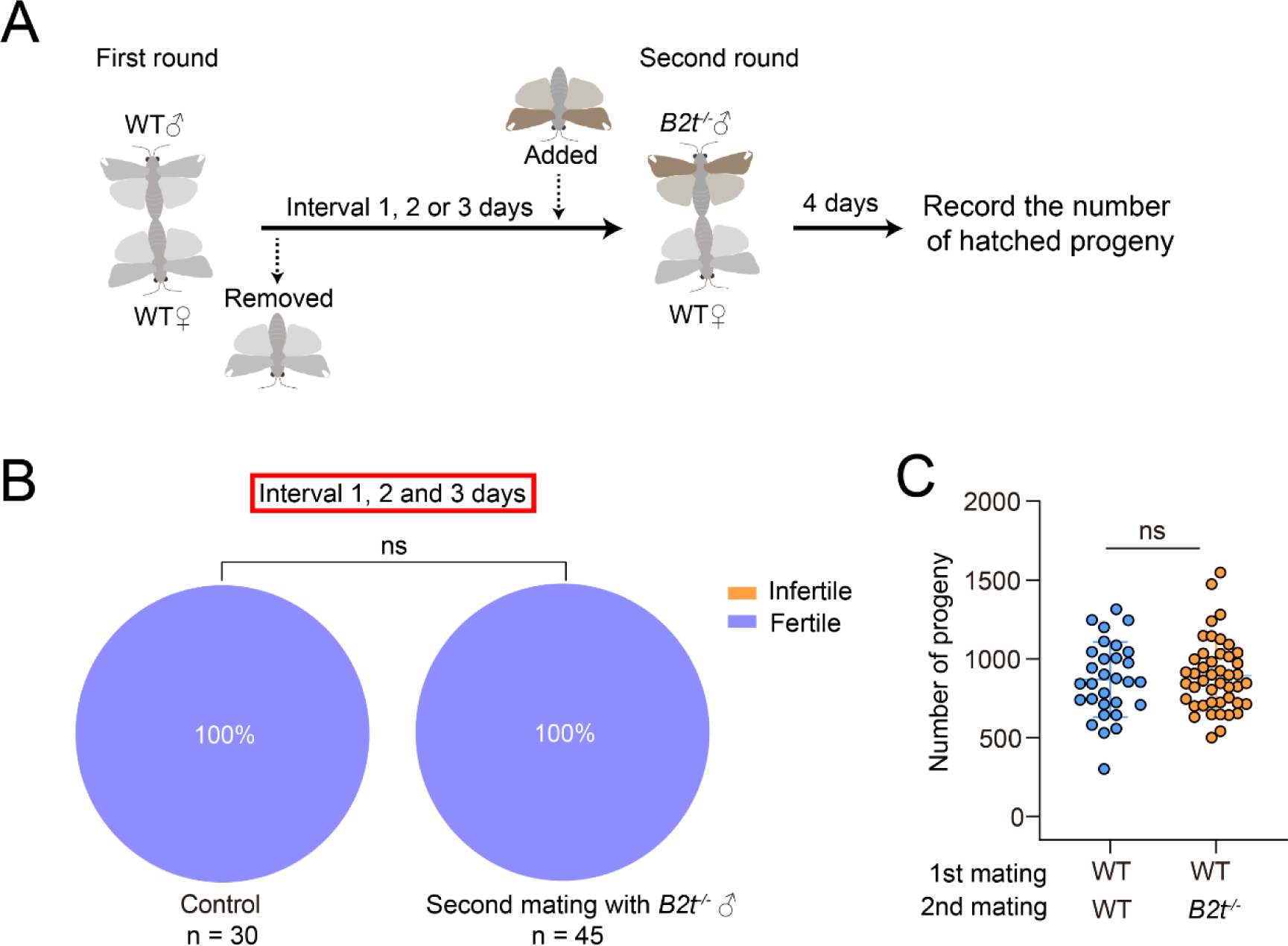
Remating with *B2t^-/-^* males did not reduce the fecundity of females pre-mated with WT males. (A) Schematic of the two rounds of single pair mating trials. (B) The percentage of fertile females from the assay (A) with different mating intervals. Fisher’s exact test; ns, not significant. (C) The number of hatched larvae from the data (B). n = 30 - 45. Error bars are mean ± SD. Student’s t test; ns, not significant.

### Release of *B2t^-/-^* males suppresses cage populations

Prior mating with *B2t*^-/-^ males could significantly suppress the fertility and fecundity of females. Therefore, we proceeded to explore whether excess release of *B2t*^-/-^ males induces population suppression. We first assessed the fitness of *B2t*^-/-^ males. Based on the comparison of life table parameters, we found no significant differences between *B2t*^-/-^ males and WT males in several parameters tested (**Figure 5-figure supplement 5A-F**). Mating competition assays showed that about 51.47% ± 7.90% of WT males and 47.30% ± 11.53% of *B2t^-/-^*males were able to successfully mate with females (marking WT, n = 3, *P* = 0.6730; marking *B2t*^-/-^ males, n = 3, *P* = 0.5969) (**Figure 5-figure supplement 5G**). The EAG recordings showed that the antennae of *B2t^-/-^*males and WT males had the same amplitude in response to stimulation with two key sex pheromones Z9-14: Ac and Z7-12: Ac (**Figure 5-figure supplement 5H**). Therefore, we predicted that *B2t^-/-^* males have comparable mating competitiveness with WT males.

Next, we conducted a cage trial to test whether *B2t^-/-^* males could suppress FAW populations when competing with WT males (**Figure 5A**). We simultaneously introduced different proportions of *B2t^-/-^* males and WT males into cages containing 15 virgin females. The number of eggs laid and hatched larvae in each cage were counted (**Figure 5B-C**). We observed that cage populations at a 1:1 release ratio had reduced offspring production compared to the control group WT release (WT only, 90.43 ± 3.03%; 1:1 ratio, 58.77 ± 9.47%) (**Figure 5C**). Moreover, the hatch rate of eggs laid in the cages declined with increasing release ratios of *B2t^-/-^* males and WT males (1:1 ratio, 58.77 ± 9.47%; 10:1 ratio, 9.58 ± 7.99%) (**Figure 5C**). Subsequently, we performed curve fitting analysis of the egg hatch rate data and found that when the number of *B2t^-/-^* males in the cage population was a multiple of 1.8 of WT males, the hatch rate was reduced by 50% compared to the control (**Figure 5D**). These results demonstrate that the release of sperm-deficient sterile males can effectively suppress FAW populations in a competitive space.

**Figure 5.**
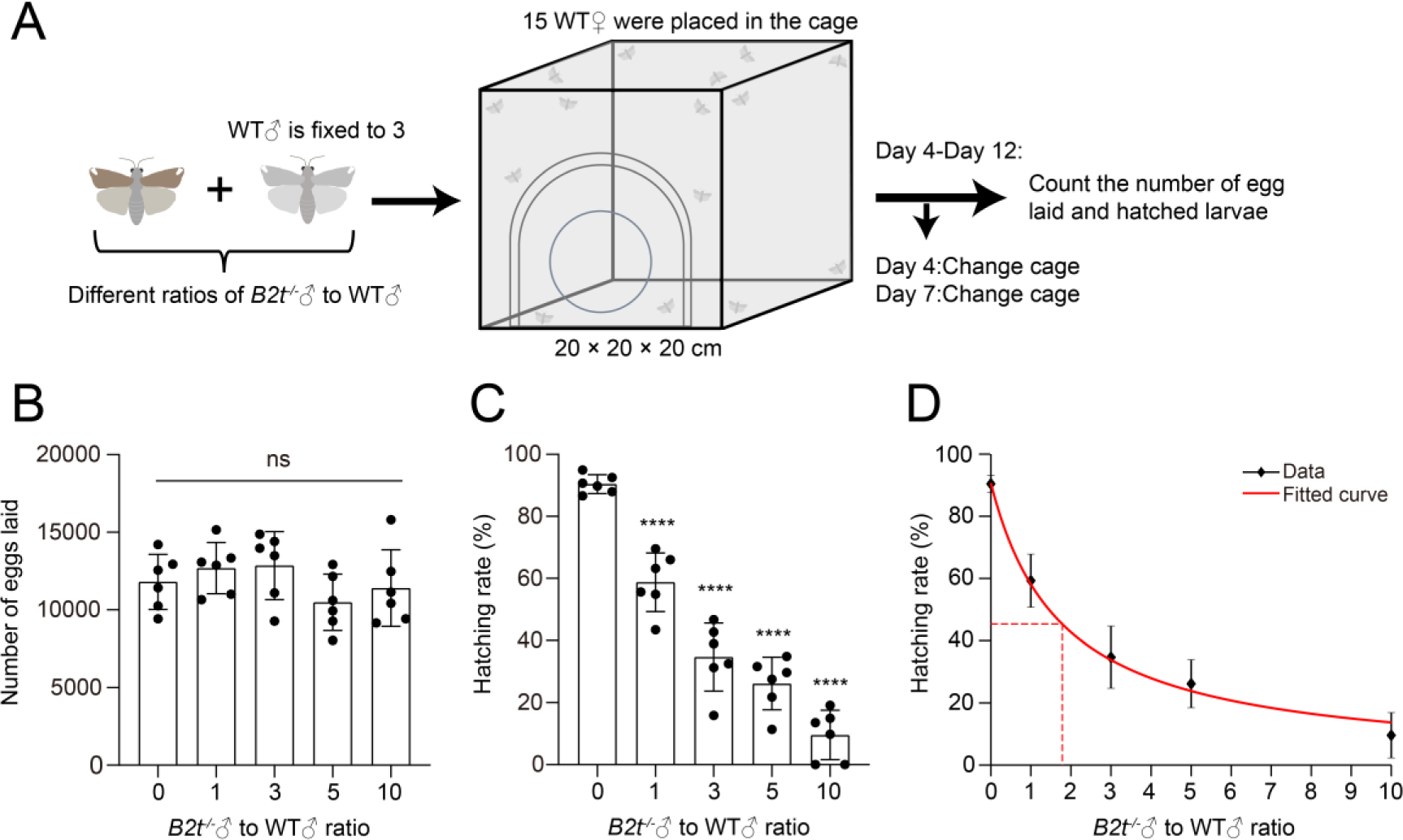
Cage release of sterile males causes population suppression. (A) Schematic of the population suppression assays performed in a 20 × 20 ×20 cm cages. (B) The number of eggs laid by females mated with different rations of *B2t^-/-^* males: WT males in each cage. n = 6. Error bars are mean ± SD. One-way ANOVA followed by Tukey’s multiple comparisons test; ns, not significant. (C) The hatch rate. Egg hatchability (%) = the number of larvae hatched / the total number of eggs laid. The hatching rate measures the extent to which the cage population is suppressed. n = 6. Error bars are mean ± SD. One-way ANOVA with Dunn’s multiple comparisons test; \*\*\*\**P* < 0.0001. (D) Curve fitting was performed on the data (C). Means ± SD. The red dashed line indicates that releasing at a ratio of *B2t^-/-^* males: WT males = 1.8: 1 reduces the cage population by 50%.

**Figure 5-supplement.**
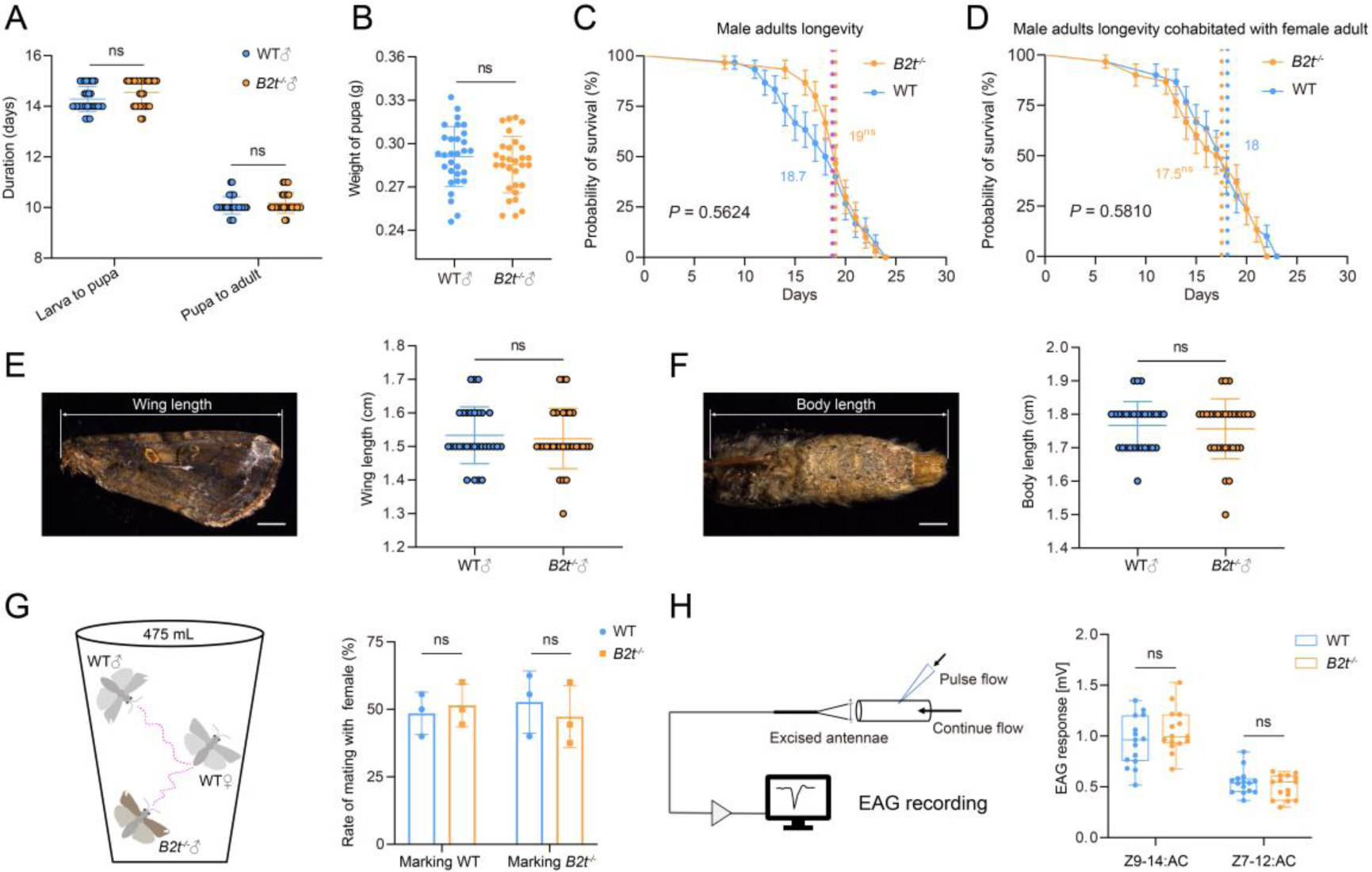
Evaluation of the fitness of *B2t^-/-^* males. (**A**) Comparison of larval and pupal periods between WT males and mutant males. n = 30 for each genotype. Mann–Whitney test; ns, not significant. (**B**) The pupal weight of WT males and *B2t^-/-^* males. n = 30 for each genotype. Student’s t test; ns, not significant. (**C and D**) Survival curves of WT males and *B2t^-/-^* males either cohabitated or isolated from WT females. Mean ± standard error (SE) is shown. Vertical dashed lines and values indicate median survivals for each test group. Statistics were performed using the Log-rank (Mantel-Cox) test, and *P* values are shown in the picture. (**E and F**) The wing and body length of WT males and *B2t^-/-^*males. The upper image shows the area used for measurement. n = 30 for each genotype. Student’s t test; ns, not significant. Scale bar, 2 mm. (**G**) The mating competitiveness of WT males and *B2t^-/-^* males. Left: Schematic of the mating competition test. Right: The rate of mating with WT female. Marking WT refers to the abdomen of WT males coated with fluorescent paint; Marking *B2t^-/-^* refers to the abdomen of *B2t^-/-^*males mounted with fluorescent paint. n = 3 groups for each treatment. Each group repeated 10 times. Student’s t test; ns, not significant. (**H**) EAG responses of excised male antennae to two sex pheromones. Left: Schematic of EAG recordings. Right: EAG responses from WT males and *B2t^-/-^* males to stimulation with Z9-14: AC and Z7-12: AC. n = 15 for each genotype. Student’s t test; ns, not significant. All data except (C and D) are presented as means ± SD.

### Modeling predicts that weekly release of sterile males will suppress FAW populations

We first conducted a paired t-test between the experimental hatch rates and the hatch rate in the simulation model under each value of *fitness*3 (see methods). Assuming that offspring viability would never be higher than females that mate with only wild-type males, we estimated *fitness*3 to be 79-100% (**Figure 6-figure supplement 6B**). Specifically, models with such a fitness range showed no significant difference (*P* > 0.01, paired t-test) between the experimental and simulated hatching rates. Thus, the sperm of WT individuals likely had an advantage relative to the sperm of *B2t*^-/-^ males because additional mating with WT males may have produced more viable offspring. When the hatching rate of females mating with one *B2t*^-/-^ male and then two or more WT males recovers to the normal level (*fitness*3 = 1, hatching rate = 91%), the simulation data is most consistent with the experimental data (**Figure 6-figure supplement 6A**). However, even this could not fully match the experimental data, indicating that the sterile males may also have had somewhat lower mating success rates in this experiment.

To explore how to eliminate the population by *B2t*^-/-^ males, we continuously every week *B2t^-/-^* males (based on a release ratio, referring to the ratio of released males to wild-type males when the population was at carrying capacity) in the model with *fitness*3 = 1. We found that when the release ratio was 3:1, the hatching rate declined to a low level, resulting in population elimination after 18 weeks (**Figure 6**). When the release ratio was 10:1, the population was eliminated in 10 weeks.

**Figure 6.**
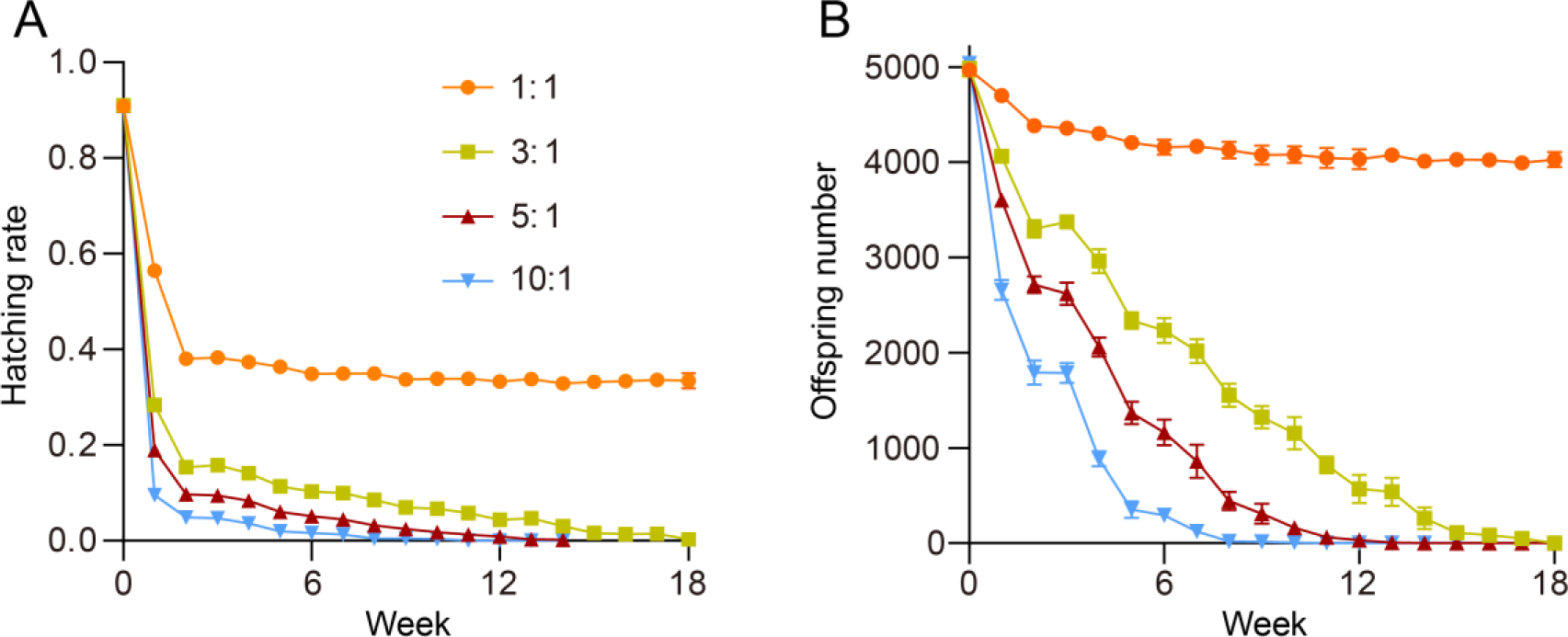
Simulation results of weekly releases of sterile males for population suppression. (**A**) The egg hatching rate through time with weekly release ratios of mutated males as shown in the figure legend. (**B**) The new viable offspring number each week. Successful population elimination occurred with a 5:1 and 10:1 release ratio. The error bars in the figure represent the standard deviation of 10 simulation replicates.

**Figure 6-supplement.**
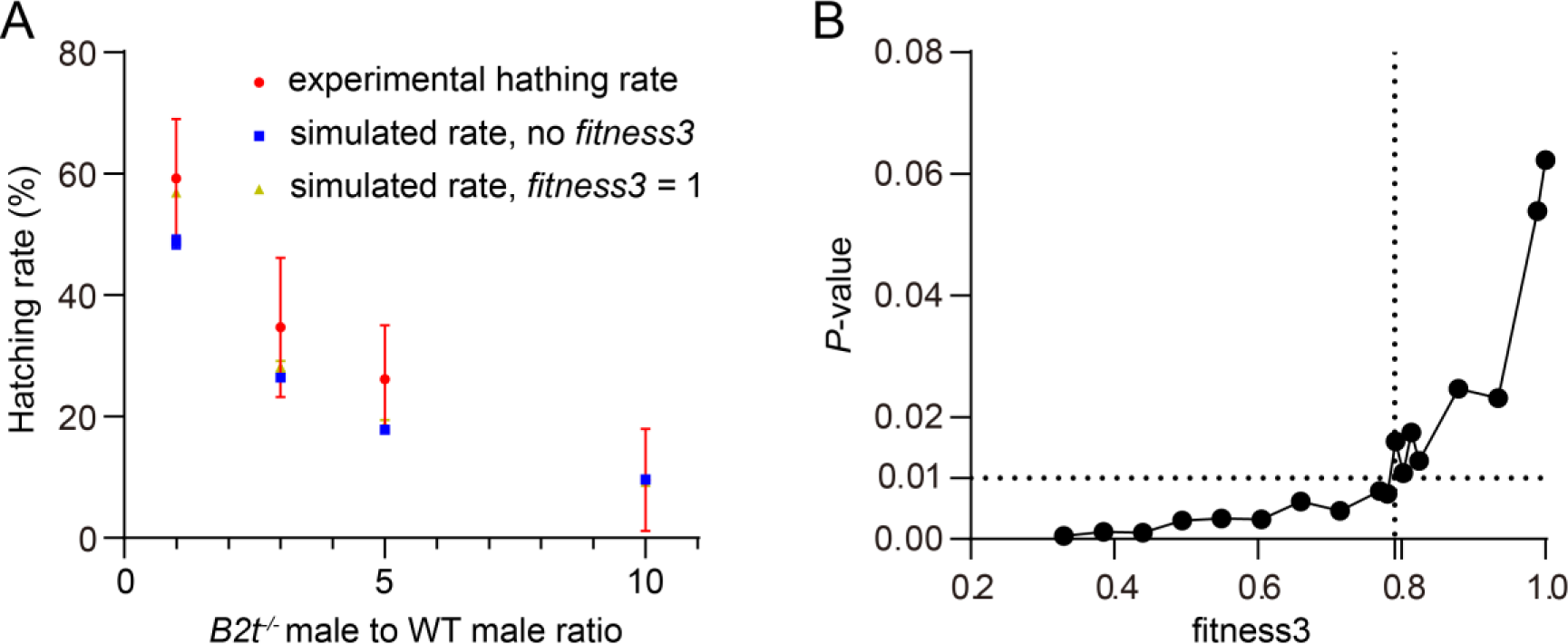
The performance of the simulation model compared to experimental results. (**A**) Shows the hatch rate tested by experiments, a simulation model in which mating with a third wild-type male has no effect after the female mates first with a sterile male and second with a WT male, and a simulation model in which *fitness*3 is 1 (offspring viability equal to wild-type after the female mates with two wild-type males after mating with a sterile male) under different release ratio. (**B**) Shows the *P*-values when the paired t-test between the experimental data and the simulation data under different values of *fitness*3.

**Table S1.**
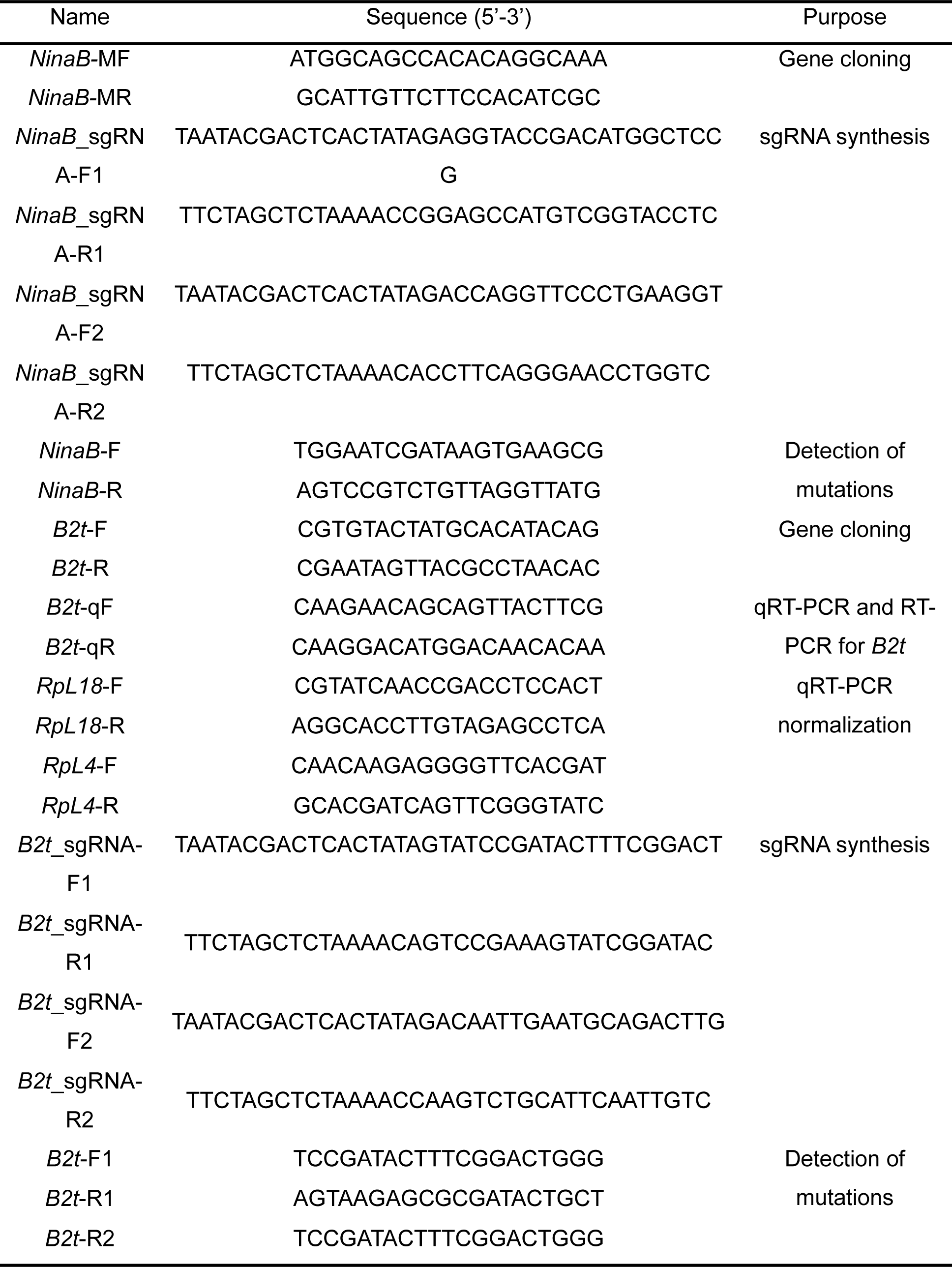
The primers used in this study.

**Table S2.**
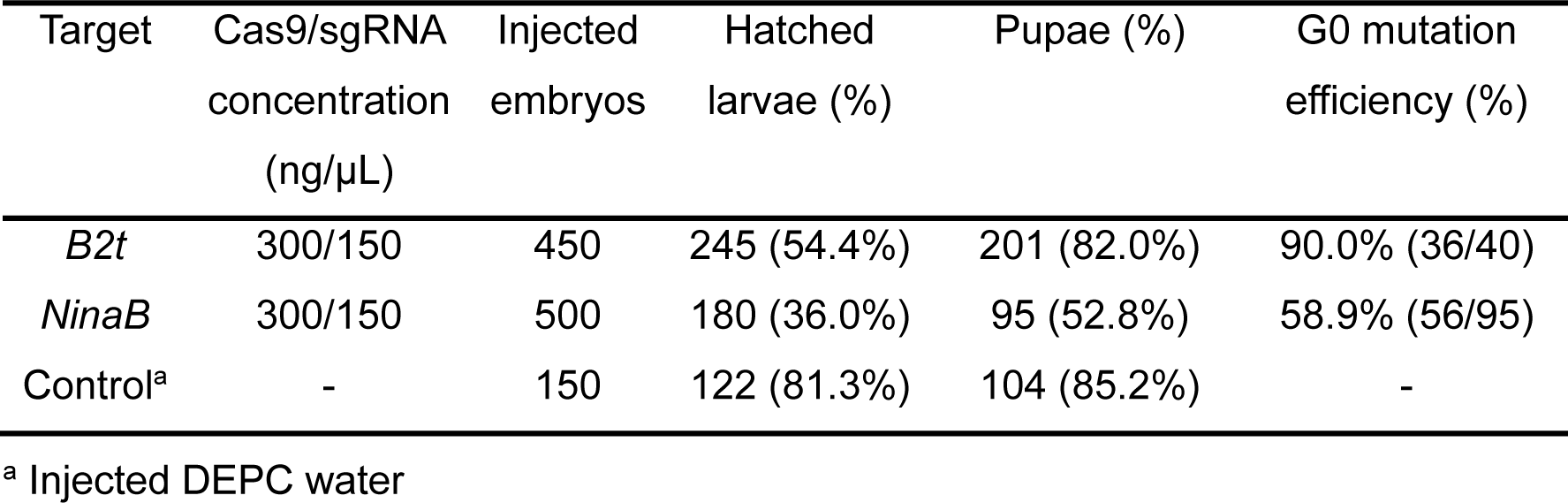
G0 mutagenesis efficiency of *B2t* induced by CRISPR/Cas9.

## Discussion

Sperm competition is an important force driving male reproductive success (1, 26). Understanding the pattern and mechanisms of sperm competition facilitates the development and application of genetic control methods. In this study, we found that FAW is a species in which females are polyandrous, but most of the offspring are sired by first males. Based on this pattern, we released sperm-deficient sterile males to successfully induce population suppression. We first found that 73% of females mated multiple times under laboratory conditions. During multiple mating, female moths had a post-mating response. Females did not gain direct benefits from multiple mating, such as increased fecundity and longevity. We used molecular markers created by CRISPR/Cas9 to clarify paternity outcomes under double mating trials with different mating intervals. Regardless of mating intervals, most females (77.3%) produced offspring using only the sperm of the first male. A few females (22.7%) also fertilized some of their oocytes with the sperm of the second male, thus increasing the genetic diversity of the offspring. Next, we constructed sperm-impaired sterile males by disrupting *B2t* gene. Mating first with *B2t^-/-^* males inhibited sperm fertilization from the second WT male, thereby significantly reducing females’ fertility and fecundity. In contrast, mating with WT before mating with *B2t^-/-^* males did not affect the number of offspring of females. Based on these results, we continued to explore the effects of releasing excess *B2t^-/-^* males on the reproduction of FAW in cages. The result showed that the release of excess *B2t^-/-^*males significantly suppressed cage population in a competitive environment. Mathematical modeling suggests that at a given population size, weekly release at high-release thresholds (*B2t^-/-^*: WT - 10:1) could completely eliminate FAW population within 10 weeks.

Multiple mattings are common in nature as a reproductive strategy for adult females (15, 27). Adult females can receive direct benefits (e.g., increased egg production and longevity) or indirect benefits (genetic benefits) from multiple mating (28–30). Several meta-analyses have shown that multiple mating of females is positively associated with reproductive output (15, 30). Significantly, the benefits to females can also drive the evolution of multiple mating (15, 31). For example, *Tuta absoluta* females derive direct benefits from multiple mating, such as increasing their egg production, fertilization rates, and longevity (32). This reproductive trait has led to the failure of control methods based on pheromone trapping of males in open greenhouses (32). We found that FAW females do not obtain gains in fecundity and longevity through multiple mating. This means that a single fertilization of females is sufficient to produce a large number of offspring. On the contrary, females have a reduced lifespan after multiple mating, which may be related to damage to females from seminal proteins, mating and egg-laying processes (33). On the other hand, genetic benefit promotes reproductive success by reducing genetic incompatibility and increasing genetic diversity (14, 28). We found that a small percentage of FAW moths could fertilize eggs with the sperm of a second male, which increases the genotypic diversity of offspring. In some social insects, research found that genetic diversity improves the fitness of offspring (15, 28, 34). Whether such an effect also exists in FAW remains to be experimentally verified.

Female adults that undergo multiple copulations provide sites for sperm competition in their reproductive tracts (35). Intense sperm competition drives the evolution of male genitalia, testes and sperm morphology to invest in sperm for reproductive success (26, 36, 37). Meta-analyses suggest that enlarged testes and longer spermatozoa are associated with adaptation to high levels of sperm competition (38, 39). In addition, mating order is often important for paternity outcomes. As in different animals such as insects and birds, the sperm of the last mating male is usually prioritized over previously intervening sperm, i.e., last-male sperm precedence (40). For example, *D. melanogaster*, *Helicoverpa armigera* and *Spodoptera litura* also use this sperm competition pattern (8, 41, 42). In contrast, we found that most FAW females (77.3%) were fertilized using only the sperm of the first male under a double mating test. Subsequently, we constructed male sterile lines with nucleated spermatozoa defects by knocking out of the *B2t* gene. Interestingly, we found that initial mating with *B2t^-/-^* males significantly suppressed the fertility and fecundity of females, even if they later mated with WT. 75% of the females that mated first with *B2t^-/-^* males could not fertilized with sperm of the second WT male to produce viable offspring. This result coincides with the identification of paternity with the help of CRISPR-based molecular markers. In Lepidoptera, apyrene sperm assist the transfer of eupyrene spermatozoa from bursa copulatrix to spermatheca (43, 44). Deletion of the *Sex-lethal* gene resulted in abnormal development of apyrene spermatozoa in the silkworm thereby causing male sterility (44). In addition, population genetic analysis of the monandrous *Manduca sexta* and the highly polyandrous *Danaus Plexippus* revealed that unfertilizing apyrene sperm proteins show little evidence for adaptive evolution in a sperm-competitive environment (25). These results suggest that apyrene spermatozoa are essential for non-competitive fertility and may not directly affect the outcome of sperm competition. Therefore, we deduce that seminal proteins may contribute in independent or synergistic patterns to the first-male sperm precedence pattern in FAW. A population genetic model examined the optimal allocation of male seminal investment at different stages of sperm competition (45). Based on the model, we considered that the first males invested seminal proteins to defend their sperm against sperm replacement by subsequently mated males. Moreover, this defense mechanism can persist for close to the entire oviposition period (3-day mating interval). However, more empirical studies are needed to validate the hypothesis that “defense” seminal proteins contributed to the first-male sperm precedence. Currently, the development of genomics and genetic manipulation tools has facilitated the study of the effects of specific seminal protein genes on paternity shares in both model and non-model organisms (12). For example, in a cricket *Teleogryllus oceanicus*, the authors found that the expression of a seminal protein gene, *gagein*, affects paternity share, and knocking down the gene resulted in a loss of paternity for the second male in the attacking position (46). Future work could further characterize the influence of one or more specific seminal protein genes on sperm competition and paternity share in FAW. Excessive introduction of *B2t^-/-^* males can suppress FAW contemporary population size. As the ratio of *B2t^-/-^* males: WT males release increased, the hatchability of eggs laid by females in the cage gradually decreased. Based on the life history and reproductive strategy of FAW, we developed a mathematical model to predict the suppression effect of weekly release of sterile males on the FAW population in the field. At release thresholds ranging from 3:1 to 10:1, the model predicts that all populations can be eliminated in 10-18 weeks. In the future, a full assessment of a release candidate would require semi-field and field trials to confirm optimal release ratios to reduce FAW populations below economic thresholds or to eliminate FAW populations. The phenomenon of female precociousness exists in the life history of FAW, with female moths eclosing on average 1.3 days earlier than male moths (Bi et al., 2023). Thus, to further improve the control level of SIT, we can release a large number of age-appropriate sterile males to fully mate first with WT females during the window period of difference in eclosion time. Further, transgenic FAW lines expressing Cas9 and gRNA were developed for gene editing (47), laying the foundation for the future establishment of pgSIT targeting *B2t* gene or other sterile male-based techniques in the FAW.

In conclusion, we performed the study from analyzing the pattern of sperm competition to managing FAW populations with sperm-deficient sterile males based on sperm competition pattern. The ejaculate of the first male, even if the nucleated spermatozoa are non-functional, can resist fertilization by the sperm of the second male. This effect can be present in females for three days. This result expands our understanding of the diversity of sperm competition patterns in Lepidoptera. We can rationalize the development of genetic control methods based on pest specific sperm competition pattern. Our results also suggest that the identification of key genes involved in apyrene spermatogenesis could serve as effective targets for FAW management.

## Materials and Methods

### Insect rearing

Each FAW larva was reared individually in a glass tube containing the artificial diet in an incubator at 26 ± 1 °C, 65 ± 5% relative humidity, and a photoperiod of 14:10 (light: dark). Pupae were sorted by gender. The newly emerged adults were transferred to 20 × 20 × 20 cm cages and fed on 10% honey water.

### Female post-mating behavior

We carried out the single-pair mating experiments to assess whether females exhibit post-mating responses. Three-day-old females were mated with a 2–3-day-old males. At 6, 24, 48, and 72 hours after first mating, each mated female was placed in a 475 ml plastic cup with a 2–3-day-old male. Successful mating rates were recorded. One 3-day-old female and one 2–3-day-old male were placed in 475 ml plastic cups for egg-laying assays. After mating, the number of eggs laid by the mated female on three consecutive days was counted. As a control, virgin females of the same age as mated females were used.

### Evaluation of the number of female copulations

To determine the number of matings of female moths throughout the adult stage, we conducted an indoor cage experiment. Fifteen female and male moths (2-3 days old) each were placed in a 20 x 20 x 20 cage to mate. Six cages were repeated, and a total of 90 females were evaluated. Survival of female moths in the cages was observed daily for the next 10 days. Dead females were immediately dissected and observed for the number of spermatophores. At the end of the 10-day period, females that were still alive were dissected and examined for the number of spermatophores.

### Multiple Mating in Females

To investigate whether multiple mating provides direct benefits to females, we compared egg production, egg hatchability and longevity of adult females after single and multiple mating. Female moths were observed every 15 minutes with a red-light source to see if they mated successfully and to record the duration of mating. For the single mating test, we removed the males after mating and left the females in the plastic cups to lay eggs. Egg production and longevity of a single female were recorded daily. For multiple mating trials, females that had mated once were allowed to mate with another virgin male. The second male was left in the plastic cup with the female (they may have mated again). We record the number of eggs laid and the lifespan of the females on a daily basis. Once the female died, we immediately isolated the spermatophores from the female’s abdomen to determine how many times the female eventually mated.

### Multiple mating in males

We conducted assays to assess the influence of multiple matings on mating behavior, fertility, and spermatophore formation in males. After a 2-3-day-old male adult mated with a 3-day-old female adult (first mating), the male was removed, and the female was left in the plastic cup. The next day, the male was mated again with a new 3-day-old virgin female (second mating). After the mating, the male was removed, and the female was left in the plastic cup. The number of eggs laid and hatched offspring were counted daily. Moths were observed every 15 min to record the female’s first and second mating duration.

The resilience of males to re-mate was tested as follows. We allowed 2-3-day-old male adults to mate with 3-day-old virgin females. At 6 and 24 hours after first mating, the male adult was placed with another 3-day-old virgin female. The mating success was determined by observations at 15-minute intervals.

For measuring the spermatophore size, the spermatophore from the copulatory bursa of the female was dissected immediately after mating and photographed using the ZEISS Stemi 2000-C microscope. The cross-sectional area of the front of the spermatophore was measured with ImageJ software.

### Gene cloning and phylogenetic analysis

With the aid of the NCBI-BLASTP program, we used the *Bombyx mori* NinaB protein and the *Drosophila* B2t protein to search for their homologs in the *S. frugiperda*. The Open Reading Frames (ORFs) were predicted using NCBI-ORF Finder. Total RNA was extracted from male adults using Trizol reagent (Invitrogen, Carlsbad, CA, USA). The complementary DNA (cDNA) was synthesized by HiFiScript cDNA Synthesis Kit (Proteinssci, Shanghai, China) according to the manufacturer’s instructions. A pair of primers were designed to clone the full-length of *NinaB* and *B2t* (**Table S1**). Alignment of the amino acid sequence was performed using ClustalX 1.81, and the comparisons were displayed using GeneDoc 2.7 software. MEGA-X was used to construct the phylogenetic tree with a neighbor-joining method.

### RT-PCR and qRT-PCR

To verify whether *B2t* is specifically expressed in the testis RT-PCR assay was conducted. The ovaries, testes, female carcass (adult without ovary), and male carcass (adult without testis) were collected on 3rd day post eclosion. RNA was extracted using Trizol (Invitrogen, Carlsbad, CA, USA). cDNAs were synthesized with HiScript II Q RT SuperMix for qPCR (+gDNA wiper) kit (Vazyme, Nanjing, China) following the manufacturer’s instructions. The cDNA and Phanta Max Master Mix (Vazyme, Nanjing, China) were used in 25 μL reaction. *RpL4* was used as a reference gene. The expression patterns of *B2t* in different developmental stages and adult tissues were studied using qRT-PCR. In addition, qRT-PCR was also used to assess the *B2t* mRNA level in *B2t* homozygous mutants (*B2t^-/-^*). RNA extraction and cDNA synthesis were the same as RT-PCR assay. The qRT-PCR analysis was performed with AceQ Universal SYBR qPCR Master Mix (Vazyme, Nanjing, China) using a QuantStudio 5 Real Time PCR System (Applied Biosystems, USA). The 20 μL reaction included 4 μL of 10-fold diluted cDNA. Three biological replicates and two technical replicates were performed for each sample. Relative mRNA levels were calculated using the 2^−△△Ct^ method. *RpL4* and *RpL18* were used as internal controls. Primers used in RT-PCR and qRT-PCR are shown in **Table S1**.

### Guide RNA design and microinjection

Two single guide RNAs (sgRNA) were designed to the exon 3 and exon 4 sequences of *NinaB*, respectively. We also designed two gRNAs to recognize the exon 2 sequences of *B2t*. sgRNAs were synthesized *in vitro* using the GeneArt™ Precision gRNA Synthesis Kit (Thermo Fisher Scientific, Vilnius, Lithuania). The Cas9 protein (TrueCut™ Cas9 Protein v2, Cat. NO. A36497) was purchased from Thermo Fisher Scientific (Shanghai, China). A mixture composed of Cas9 protein (300 ng/μL) and sgRNA (150 ng/μL) was co-injected into embryos (<1 hr-old) using the InjectMan NI 2 microinjection system (Eppendorf, Hamburg, Germany). Injected eggs were incubated at 26 ± 1 °C and 70% ± 5% relative humidity for hatching.

### Generation and characterization of *NinaB* and *B2t* mutants

Mutation detection and crosses were performed following the established method (48). Sequencing of the G0 mutants showed multiple peaks near the target site, suggesting mutations. The PCR products were cloned into the pClone007 Simple vector (TSINGKE, Beijing, China) and sequenced. The G0 mutants were mated with the wild-type (WT) in single pairs. After mating and laying eggs, G1 eggs were genotyped. The type of mutation inherited from the G0 was identified. Single-pair sibling crosses of G1 mutants produced G2 progeny. For the construction of *NinaB* mutants (*NinaB^-/-^*), we chose a pair of primers F/R to amplify the target region. DNA amplification products were sent for sequencing to identify the *NinaB*-null allele. For generating *B2t* mutants, the forward primer F1 and the reverse primer R1 were designed to detect the *B2t* mutation (Fig. 2A). A reverse primer R2 located at the target site 1 was used to distinguish *B2t^-/-^* homozygous mutants from *B2t* heterozygotes (*B2t^+/-^*) based on the band pattern of PCR amplified products.

We conducted fecundity analysis assay to investigate whether *NinaB* and *B2t* mutation impairs male fertility. For determining the fertility of *NinaB^-/-^*males, 12 WT or *NinaB^-/-^* males were mated with WT females. To explore whether *B2t* mutation affects male fertility, we used five different mating combinations: 40 pairs of WT males + WT females, 40 pairs of WT males + *B2t^-/-^*females, 40 pairs of *B2t^+/-^* males+ WT females, 40 pairs of *B2t^-/-^* males+ WT females, and 40 pairs of *B2t^-/-^* males + *B2t^-/-^* females. A pair of 1-2 days old adults were transferred to 475 mL plastic cups for mating. When the female adult began laying eggs, we recorded the number of eggs laid for four consecutive days. Finally, the total number of eggs laid and hatched larvae were recorded.

### Analyzing competitive fertilization results

Based on the result that the knockout of *NinaB* does not reduce male fertility, we selected the *NinaB* mutation as a molecular marker. We conducted a double mating trial with different mating intervals to determine paternity. WT females were first mated to WT males. One, two or three days after the first mating, females were successfully mated with *NinaB^-/-^* males. We daily recorded the number of eggs laid by female adults. In addition, we extracted DNA from all eggs laid by female adults after completion of the second mating. A pair of primers F/R was used for PCR to identify the presence of *NinaB* mutant allele in the progeny. *NinaB* mutant males used for mating in the experiment were of the same age (3 old days) and body weight as WT males.

### Fluorescence staining

Fluorescence staining of sperm bundles was performed as previously described (49). Testes were dissected from 1-day-old WT males and *B2t^-/-^* males. Sperm bundles were collected in 1.5 mL centrifuge tubes and fixed in 4% paraformaldehyde for 1 hour. At the end of fixation, the samples were washed three times with PBS. Subsequently, the tissues were incubated with TRITC Phalloidin (YEASEN Cat#40734ES75, 1:1000) for 1 hour and DAPI (Solarbio Cat#C0060, 1:1000) for 10 minutes. After staining, the samples were washed three times with PBS. The tissues were imaged using a ZEISS LSM 980 with Airyscan 2, and the pictures were processed with ImageJ software.

### Effect of prior mating with *B2t^-/-^* males

To determine whether WT females mated for the first time with *B2t^-/-^* males had a post-mating response. We conducted a two-round mating experiment. WT female first mated with *B2t^-/-^* males. After observing the end of mating, *B2t^-/-^*male was removed. At 24h,48h or 72h after the first mating, WT males were added to court and mate with WT females that had mated once. After the second mating was completed, the WT males were removed. The proportion of females that received virgin WT males was recorded. Females first mated with WT as a control group.

Meanwhile, double mating trials with different mating intervals were also carried out to investigate whether prior mating with *B2t^-/-^*males could affect competitive fertilization. The number of eggs laid and hatched was recorded. The copulatory bursa from the female was dissected to examine the spermatophores. The presence of two spermatophores shows that the female had completed the second round of mating. In control experiments, female adults were mated with WT males in both rounds.

### Effect of second mating with *B2t^-/-^* males

To explore whether the second mating of females to *B2t^-/-^*males affect paternity, we conducted an additional two rounds of single-pair mating trials. The experimental methods used are the same as those described above, except that the mating order of females was switched: females were first mated with WT males and then with *B2t^-/-^* males.

### Fitness evaluation

Life tables were established to compare the fitness parameters between *B2t^-/-^* males and WT males. Each larva was individually reared in a glass tube (25 × 75 mm) with the artificial diet until pupation. During the early larval stages, larvae were observed every two days for feeding status. At the end of the sixth instar, the larvae were observed twice daily to record pupation. One day after pupation, male pupae were identified based on morphological markers described previously (50) and placed in a plastic tube (25 × 95 mm). *B2t^-/-^* pupae males were identified by PCR-based genotyping using F1 and R2. The larval and pupal development time of WT males and *B2t^-/-^* males were recorded. Survival curves were prepared to record the lifespan of male adults cohabitating and non-cohabitating with female adults. Newly eclosed male adults were placed by themselves or cohabitated with a newly eclosed WT female in a 475 mL plastic cup. The moths were fed on 10% honey water. Mortality was recorded daily until all males died. The 2-3-day-old male adults were anesthetized with carbon dioxide, and the body and wing length of WT males and *B2t^-/-^* males were determined using vernier scale.

### Assessment of male mating competitiveness

To estimate the mating competitiveness of *B2t^-/-^* males, 2-3 days-old *B2t^-/-^* males and WT males were mated with 3 days-old WT females in 475 mL plastic cups. The abdomens of males were coated with a green fluorescent paint to mark the genotype and the mating was recorded at 15 min intervals. Once we found that mating was occurring, we illuminated the abdomen of the male adult with a violet light source to check which genotype of successful mating male. To eliminate the possible effect of fluorescent paint on mating, we carried out two sets of experiments in parallel. WT males were coated with fluorescent paint in one experiment, and *B2t^-/-^*males were coated with fluorescent paint in the other experiment.

### Electroantennogram recording analysis

Electroantennogram (EAG) recordings were carried out to measure the response of WT males and *B2t^-/-^* males to two sex pheromone (Z)-9-tetradecenal acetate (Z9-14: Ac) and (Z)-7-dodecenyl acetate (Z7-12: Ac) (purity >92%, Kunming Bohong Biotechnology Co., Ltd). Two sex pheromones were dissolved in hexane at a final concentration of 100 ng/μL and hexane was used as a control. The excised antennae of 3-day-old virgin males were used in EAG recordings. The EAG recordings test was performed with reference to the previously reported methods (51). Briefly, we loaded 10 μL test solution onto a 2 × 1 cm filter paper. After the filter paper evaporated for 5 minutes, the filter paper was placed in a Pasteur pipette, and we closed the mouth of the tube with a parafilm (Bemis, USA). The test antenna was connected by a conductive gel (SPECTRA 360, Fairfield, NJ, USA) to the Y-type conductivity meter. Then, the conductivity meter and Pasteur pipette were placed into the EAG instrument (Syntech, Germany). The pulse duration (4 mL/s) of the odor stimulus was tested at 0.5 s. After 30 seconds of baseline stabilization, the excised antennal responses to sex pheromone and hexane were tested. The interval between stimuli was greater than 50 s to recover the response sensitivity of the antennae. EAG values were recorded and analyzed using the EAGPro software (Syntech, Germany).

### Cage release assay

We conducted a population suppression experiment to investigate whether the release of *B2t^-/-^* males in a scenario with WT males could suppress FAW populations. We simultaneously released 2-3 days old *B2t^-/-^*males and WT males in 4 different ratios (*B2t^-/-^* males: WT males = 1, 3, 5, and 10) into a 20 × 20 ×20 cm cage containing 15 2-3 days old females. The group with a 0:1 ratio of *B2t^-/-^* males to WT males was used as a control. Fresh 10% honey water was used as a daily source of nutrition. The number of WT males was fixed at 3. The test adults were transferred to new cages on days 4 and 7. The number of eggs laid and hatched larvae were recorded over 12 days (Day 4 - Day 12) starting from the day of egg laying.

### Curve Fitting

Curve fitting of egg hatch data from cage release assay data were performed as previously described (24). The formula: P = P0 / (1 + K*X) × 100% was used. P0 is the maximum hatching rate of WT females in the absence of *B2t^-/-^* males’ release. K is the competitiveness of *B2t^-/-^*males normalized to WT males. X is the release ratio of *B2t^-/-^*males to WT males. Based on the experimental data, the parameters P0 and K were calculated using the maximum likelihood estimation. P0 = 0.9071 ± 0.0021 (mean ± 95% C.I.); K = 0.5615 ± 0.0055 (mean ± 95% C.I.).

### Mathematical modeling

According to the life history of *Spodoptera frugiperda*, we constructed an individual-based panmictic model. In this model, the lifespan of individuals is four weeks. The first two weeks are larva, and the last two weeks are adults. Only adults can mate and reproduce. Each 3-week-old female randomly selects a mate, and the probability of successful mating and reproduction is equal to the fitness of the chosen male. 78% of females can remate in the first week, and 53% of females can mate three or more times before offspring are produced. The number of fertilized eggs follows normal distribution with mean of 1000 and standard deviation of 242. Because about 9 percent of eggs cannot successfully be hatched in a natural environment, the hatching rate when the first mate is wild-type follows a normal distribution with mean of 0.91 and standard deviation of 0.0303, according to experimental data. The hatching rate of females mating only with sterile mutant males is 0. If a female mates with a wild-type male after mating with a sterile mutant male in the third week, then there is a 75% chance that the offspring hatching rate is 0. For the remaining 25 percent, the hatch rate is drawn from a normal distribution with mean of 0.47 and standard deviation of 0.19. Because 53% percent of females mate three times or more (experimental paper citation), we modified some of our simulations. We did not determine the experimental hatching rate in this case, but we instead assume that if a female first mates with a mutated sterile male and then mates with two or more wild-type males, the hatching rate of this female would recover to *fitness*3 compared to females that mate only with wild-type (thus giving a 91% hatch rate if *fitness*3 = 1). This increased fitness could be due to a sperm advantage of wild-type males. Here, we varied the value of *fitness*3 to find the range of its real value.

Because the environment and resources limit larval survival, they compete with each other after reproduction. Offspring survival rate is calculated as:

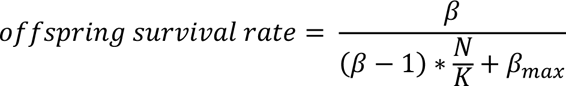

where β represents the low-density growth rate of the population, *N* represents the number of newborn offspring, *K* represents the environmental carrying capacity of adult females in a normal wild-type population, and β_*max*_ represents the maximum number of offspring from each female. This formula simulates a Beverton–Holt density-dependent growth curve to keep the population stable (52).

All the simulations are implemented in the SLiM forward-in-time population genetic simulation framework, version 4.0.1 (53). We set the capacity as 20,000, allowed 10 weeks for the natural population to equilibrate, and then released different ratios of sterile mutant males each week (the ratio refers to the number of released males for each wild-type male in a wild-type population at carrying capacity). We recorded the hatching rate of all females and the survival of offspring in each week. We used 10 simulation replicates for each release ratio and used mean square error to evaluate the model.

### Statistics

The GraphPad Prism 8 software package (GraphPad, San Diego, CA, USA) was used for statistical analysis and graphing of the data. The data were first verified for normal distribution by the D’Agostino–Pearson normality test. When data were normally distributed, parametric tests were used; when data were not normally distributed, we used non-parametric tests. The sample sizes and statistical tests for each experiment are shown in the Figures or described in Figure legends.

## Acknowledgments

This research was funded by the National Key R&D Program of China to SFW (2022YFD1700200) (https://service.most.gov.cn/) and a National Natural Science Foundation of China grant to SFW (32022011) (https://www.nsfc.gov.cn/).

The funders had no role in study design, data collection and analysis, decision to publish, or preparation of the manuscript.

## Declarations

### Ethics approval and consent to participate

Not applicable.

### Consent for publication

Not applicable.

